# Adapting Co-Folding Models for Structure-Based Protein-Protein Docking Through Flow Matching

**DOI:** 10.1101/2025.11.28.691195

**Authors:** Da Xu, Lee-Shin Chu, Jeffrey J. Gray

**Affiliations:** Department of Chemical and Biomolecular Engineering, Johns Hopkins University, Baltimore, MD, USA

## Abstract

Co-folding models like AlphaFold have revolutionized protein complex structure prediction, yet their reliance on multiple sequence alignments (MSAs) limits their applicability on challenging targets such as antibody-antigen complexes. An alternative approach, structure-based protein-protein docking, predicts the bound complex structure from the unbound monomer structures without requiring MSAs. In this work, we propose a novel method to adapt co-folding models for structure-based protein-protein docking by replacing their template module with a docking module, followed by training end-to-end with a flow-matching objective. We apply our method to AlphaFold-Multimer (AF-M) using the OpenFold implementation and transform it into a generative docking model, which we name AF2Dock. We evaluate AF2Dock and various baseline methods on the PINDER-AF2 benchmark and an antibody/nanobody test set. When using non-holo inputs, AF2Dock shows competitive performance compared to other structure-based docking methods and, in the case of nanobody complexes, outperforms all other docking methods tested here. Although AF2Dock underperforms co-folding AF-M and AF3 in success rates when using non-holo inputs, it produces orthogonal predictions and successfully identifies correct structures for targets where co-folding models fail. Ablation studies confirm that full-parameter fine-tuning of the AF-M components is critical for performance and reveal that, surprisingly, the inclusion of ESM embeddings can hinder success rates in certain cases such as nanobody complexes. The code is available at https://github.com/Graylab/AF2Dock.

## I. INTRODUCTION

Protein-protein interactions mediate critical processes in all forms of life, and the accurate prediction of the three-dimensional details of such interactions can provide significant insights into many biological questions. The current state-of-the-art models for predicting protein complex structures are co-folding models such as AlphaFold and related models [1–6], which infer inter-residue contacts primarily using sequence information by extracting evolutionary signals from multiple sequence alignments (MSAs) of related proteins. Although many co-folding models also allow the input of structural information through templates, such information is found to have only minor effects on the prediction unless the signal from the input MSA is weak [7, 8].

Despite increased prediction accuracy of protein complexes brought about by co-folding models, performance on this task still lags behind that of single-chain protein structure predictions [2]. This discrepancy may be explained by the stronger evolutionary constraints within a single protein chain, resulting in richer co-evolutionary signals on intra-chain contacts in MSAs, compared to the often weaker and noisier signals of inter-chain contacts. There have also been observations that suggest a non-intuitive role of MSAs beyond inter-chain co-evolution in co-folding models for protein complex predictions. For instance, using unpaired multi-chain MSAs (where chains in each row of the concatenated MSA can come from different organisms) can still lead to comparable prediction accuracy [9], and models like Chai-1 can achieve good complex prediction accuracy by using only ESM embeddings [3], which only contain information akin to single-chain MSAs [10]. Although the mechanisms behind how co-folding models utilize MSAs for inter-chain prediction require further exploration, the reliance on MSAs could be a limiting factor for these models, especially in cases where there is low availability of related sequences or when the evolutionary process is unique (such as antibody-antigen or pMHC-TCR complexes).

In contrast to co-folding models, a more traditional approach for modeling protein-protein interactions is structure-based protein docking, which aims to predict bound protein complexes given the unbound monomer structures [11]. Instead of utilizing evolutionary information, structure-based protein docking methods infer interactions based on structural features such as shape complementarity, surface properties, and residue-residue interactions. For both physics-based and deep-learning-based protein docking methods, a common strategy is to split the docking process into two parts: initial generation of many possible structures with a sampling model, followed by ranking of those generated structures based on metrics computed from the structures. In the case of deep-learning-based methods, diffusion-based generative docking models have recently emerged as a promising direction due to their close alignment with the sample-and-rank scheme as well as their ability to model complex distributions [12–15]. DiffDock-PP [12] is the first model to demonstrate this process, with separate sampling and ranking models. DFMDock [13] unifies sampling and ranking into a single model, and shows better generalization compared to DiffDock-PP. Nevertheless, current diffusion-based docking models are all limited to rigid-body docking and considerably underperform co-folding models due to their inability to properly rank the generated structures.

Although trained as a co-folding model dependent on MSAs, AlphaFold has been shown to have learned an energy function that is independent of evolutionary information. In the case of AlphaFold2 (AF2), Roney and Ovchinnikov [16] demonstrated this through the AF2Rank approach, where protein structures of different quality are input as templates into AF2 in single-sequence mode without MSAs. They observed that the confidence scores output by AF2 strongly correlate with the quality of the input structures. In the case of AlphaFold-Multimer (AF-M), Mirabello et. al. [17] performed similar experiments with full-complex templates and observed the same correlation between confidence score and prediction quality. Interestingly, they also observed that in certain cases, when inputting the ground truth template, using MSAs resulted in worse predictions compared to single-sequence mode [17], providing evidence that spurious signals in MSAs could impair performance.

An interesting research question is whether one could harness the intrinsic energy function of AlphaFold for structure-based protein-protein docking without using MSAs. A naive approach is to use AlphaFold in single-sequence mode with custom template inputs, where only the protein sequences of individual subunits are input into the model along with their structures as templates to predict the complex structure without MSAs. Anecdotally, this approach has proven successful in certain cases [18], but a comprehensive evaluation of its performance is not available. In this work, we develop an alternative approach for adapting AlphaFold as a generative model for structure-based protein-protein docking by replacing its template module with a novel docking module and training end-to-end with a flow-matching objective [19]. We test our approach on AF-M [1] using the OpenFold implementation [20], and we train with the PINDER training set [21]. We name the resulting model AF2Dock and evaluate its performance against various baseline methods on the PINDER-AF2 benchmark [21] and an antibody/nanobody test set [22]. When using non-holo structure inputs, we show that AF2Dock performs competitively with baseline docking methods and, in the case of nanobody complexes, outperforms all other docking methods tested here. Additionally, we show that although AF2Dock underperforms co-folding AF-M and AF3 in success rates when using non-holo inputs, it produces orthogonal predictions and successfully identifies correct structures for targets where co-folding models fail, which could allow better success when the results from the two methods are combined.

## II. METHODS

Fig. 1 shows the schematics for the AF2Dock inference process and its model architecture. AF2Dock is akin to AF-M in single-sequence mode, where single protein sequences are used as input instead of MSAs. Given a noisy protein complex structure, AF2Dock is tasked with denoising it into the bound complex structure. We train AF2Dock with a flow-matching objective, and its inference involves an iterative process that integrates along the learned velocity field **v**_*t*_(**x**_*t*_; *θ*) with a set of intermediate time points *t* from *t* = 0 (prior distribution *p*_0_) to *t* = 1 (data distribution *p*_1_) (Fig. 1a). As a generative model, we sample many structures for each target from different initial noises, and we rank the resulting predictions with AlphaFold’s confidence score ipTM. Although here we apply our methodology to AF-M, this scheme can be extended to other AlphaFold-like models.

**FIG. 1.**
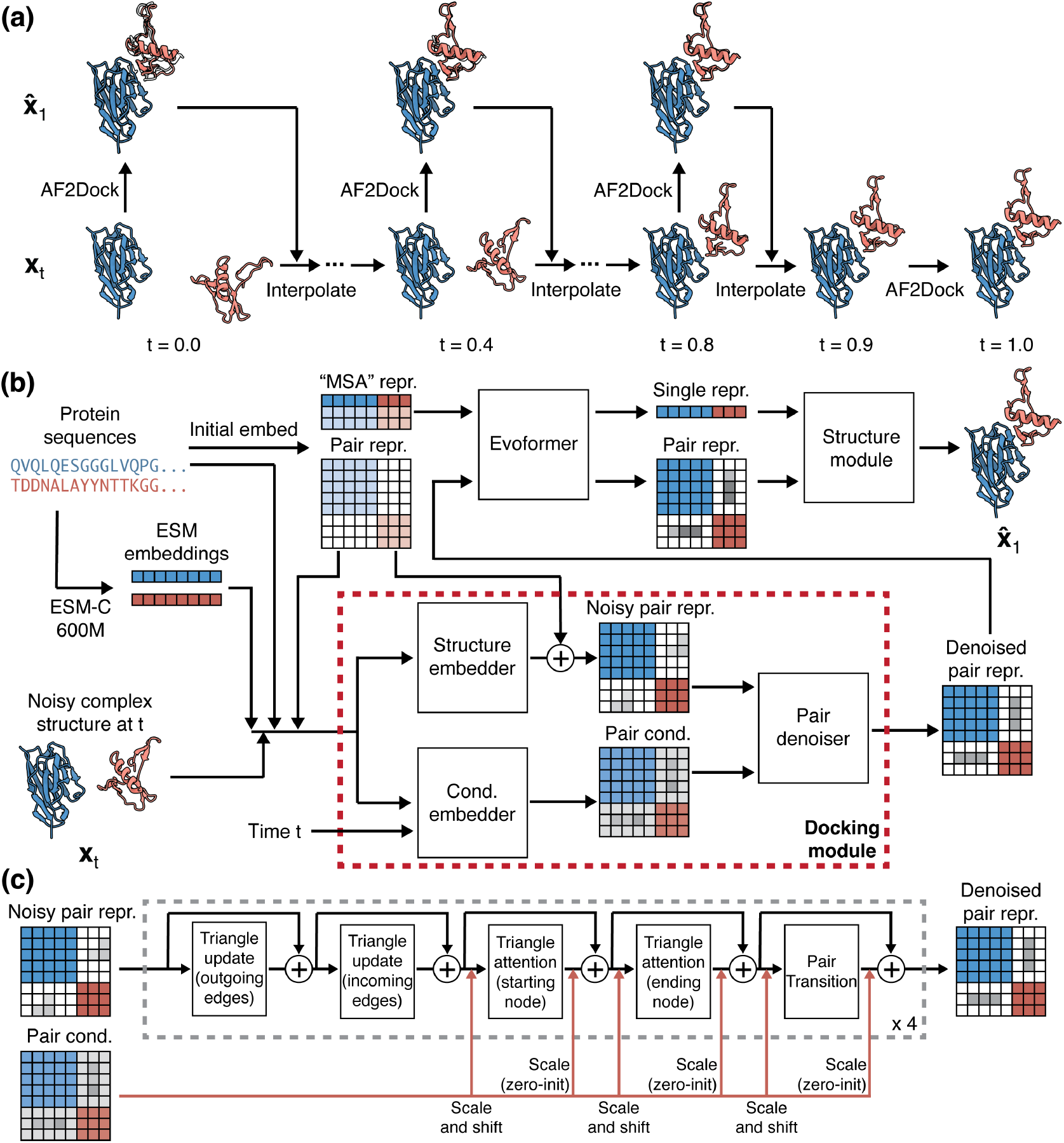
Schematics for AF2Dock inference and architecture. (a) Inference process for AF2Dock. Starting from a noisy structure **x**_0_ sampled from the prior distribution *p*_0_ at *t* = 0, the structure is iteratively denoised through a set of intermediate time points *t* to produce **x**_1_ in the data distribution *p*_1_ at *t* = 1. At each time point *t*, AF2Dock is used to predict the fully denoised 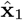 given **x**_*t*_, which are then interpolated to produce **x** at the next time point. The final predicted **x**_1_ is shown along with 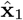 as silhouettes. Nanobody-bound SARS-CoV-2 Nsp9 (PDB: 8dqu) is used for illustration. (b) Architecture of AF2Dock. AF2Dock is akin to AF-M in single-sequence mode, where the MSA representation is produced from single protein sequences (but not the ESM embeddings). The template module of AF-M is replaced with a docking module that takes inputs of noisy complex structure **x**_*t*_ along with sequence information to predict a denoised pair representation. This denoised pair representation is fed back into the Evoformer to complete the folding pipeline. (c) Schematics for the pair denoiser module. In each block, the noisy pair representation is processed through a pair stack, including triangle multiplicative update, triangle attention, and pair transition. The pair conditioning influences the denoising process through adaptive layer normalization before and after the triangle attention and the pair transition blocks. We use a total of four blocks for the pair denoiser module.

### A. Docking module

The key architectural innovation of AF2Dock is the replacement of the template module in AlphaFold with a docking module (Fig. 1b). In contrast to the template module, which only performs embedding of the input structural information, our docking module performs both embedding and denoising to produce a denoised pair representation based on the noisy structure input **x**_*t*_.

The noisy structure input **x**_*t*_ is first embedded into a noisy pair representation by the structure embedder through a similar process as the template module [1, 2] (Algorithm S1). We convert the structure input into the same set of pair features as used by the template module, including the distogram (computed from pair-wise distances of C_*β*_ atoms), unit vectors (computed from pair displacement vectors of backbone frames), as well as C_*β*_ atom masks and backbone frame masks. Similar to AlphaFold-Unmasked [17], we do not mask out the inter-chain information as AF-M does [1]. In addition to structural features, we embed positional features including amino acid identities as well as the ESM-C embeddings [23] computed from individual protein sequences. The features are combined and processed through a two-block pair stack [1, 2], same as in the template module.

To enable denoising of the noisy pair representation, we adopt a diffusion-transformer-like architecture [24] used in many diffusion models including AlphaFold 3 (AF3) [2]. This involves a pair conditioning embedding process and a conditioned denoising process, performed by the conditioning embedder and the pair denoiser. The pair conditioning is generated using the same set of features as the noisy pair representation, with the addition of the time conditioning (Algorithm S2). The pair denoiser (Fig. 1c) then takes input of the pair conditioning and the noisy pair representation to perform conditioned denoising (Algorithm S3). In each block of pair denoiser, the noisy pair representation is processed through a pair stack conditioned using the adaptive layer normalization (AdaLN) approach from the diffusion transformer [2, 24]. Specifically, we use the pair conditioning to predict the element-wise scaling and shifting parameters, which are applied to the pair representation before and after the triangle attention and pair transition steps.

The output denoised pair representation is fed back into the Evoformer to complete the rest of the folding pipeline. We did not modify the architecture of AF-M outside of the template module, except that we disabled recycling. The reasoning for disabling recycling is two-fold. First, if we perform recycling at each time step, the total number of network evaluations would be the number of time steps multiplied by the number of recycles, which can make the runtime impractical for large complexes. Second, we theorized that the iterative denoising process of flow-matching functions similarly to recycling, where the network is evaluated multiple times for each sample, and adding recycling on top of that may not be necessary. For the docking module, an intuitive perspective to understand its role is that it provides AF-M with an initial guess by denoising the noisy input complex structure, and this initial guess is then refined through Evoformer to produce the final prediction.

### B. Flow-matching objective

Flow matching (FM) is a generative modeling paradigm [19, 25, 26] that, given data distribution *p*_1_ and a prior distribution *p*_0_, defines a probability path *p*_*t*_(**x**_*t*_) that transforms *p*_0_ to *p*_1_. The time *t* here is an abstract dimension that describes the degree of transition between the two distributions and is not physically meaningful. The probability path *p*_*t*_(**x**_*t*_) has a corresponding velocity field **u**_*t*_(**x**_*t*_) that governs the evolution of **x**_*t*_ through an ordinary differential equation (ODE):

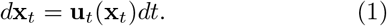

The FM objective aims to learn the velocity field **u**_*t*_(**x**_*t*_) using a neural network **v**_*t*_(**x**_*t*_; *θ*) parametrized by *θ* [19]:

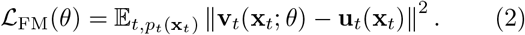

As an explicit definition of data distribution *p*_1_ (and potentially the prior distribution *p*_0_) is typically unavailable, explicitly defining the probability path *p*_*t*_(**x**_*t*_) is intractable. Instead, the neural network is trained with a conditional flow matching (CFM) objective [19]:

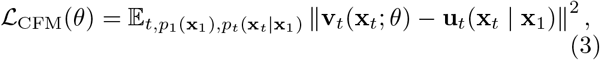

where **u**_*t*_(**x**_*t*_|**x**_1_) is the conditional velocity field corresponding to a conditional probability path *p*_*t*_(**x**_*t*_|**x**_1_), both conditioned on a single data point **x**_1_. These conditional quantities can recover the original (marginal) probability path *p*_*t*_(**x**_*t*_) and velocity field **u**_*t*_(**x**_*t*_) through marginalization over the data distribution *p*_1_:

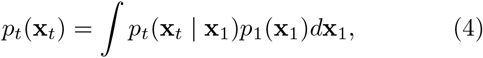

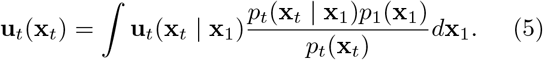

In contrast to the marginal probability path *p*_*t*_(**x**_*t*_), the conditional probability path *p*_*t*_(**x**_*t*_|**x**_1_) is easy to define as it only depends on a single data point **x**_1_ and can be chosen as desired. Optimizing the CFM objective (Eq. 3) is shown to result in identical gradients with regard to neural network parameters *θ* as the FM objective (Eq. 2) [19]. This means that although we match the conditional velocity field **u**_*t*_(**x**_*t*_|**x**_1_) during training, the model is still learning the marginal velocity field **u**_*t*_(**x**_*t*_) implicitly defined by our chosen **u**_*t*_(**x**_*t*_|**x**_1_).

The CFM objective has been previously employed to adapt AF2 for predicting conformational ensembles of proteins [27]. Here, we use the CFM objective to train AF2Dock for generative structure-based protein-protein docking. Given a single data sample **x**_1_ and noise **x**_0_ sampled from the prior distribution *p*_0_, we define the conditional probability path as the conditional optimal transport path [19, 27]

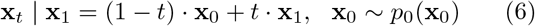

with a corresponding conditional velocity field

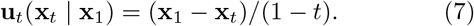

As in [27], instead of directly learning a velocity field **v**_*t*_(**x**_*t*_; *θ*), we define our neural network **x**(**x**_*t*_, *t*; *θ*) via reparametrization

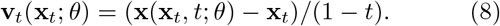

Substituting Eq. 7 and 8 into Eq. 3 results in the following objective:

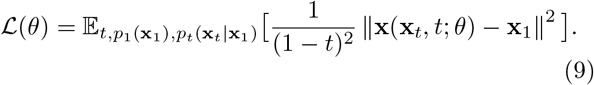

Thus, our neural network learns to predict the expectation of the fully denoised sample 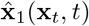 given inputs **x**_*t*_ and *t*:

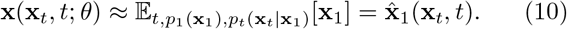

Due to the *SE*(3)-equivariant nature of AF-M, instead of directly using the mean squared error loss as shown in Eq. 9, we use the original AF-M losses [1] for training (with the omission of the masked MSA loss):

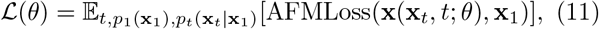

which allows not only training AF2Dock’s ability to predict correct structures with the frame-aligned point error (FAPE) loss but also aligning its confidence output such as the ipTM score with the prediction quality through auxiliary losses.

We define the prior distribution *p*_0_ as the joint distribution of the rigid-body prior *p*_0,rb_ and the internal flexibility prior *p*_0,flex_. The rigid-body prior *p*_0,rb_ is defined over the rigid-body displacement of the ligand protein with regard to the receptor protein, and can be decomposed into translational and rotational components in ℝ^3^ and *SO*(3). We define the prior distribution for translation as a Gaussian distribution 𝒩 (0, *σ*_tr_*I*_3_) and as a uniform distribution 𝒰 (*SO*(3)) for rotation. For the internal flexibility prior *p*_0,flex_, here we consider only small perturbations defined as the collection of all realistic unbound protein structures. Practically during training, we approximate this distribution by sampling from all available unbound monomer structures in the PINDER training set [21] (including holo, apo, and predicted monomer structures). A potential future direction is to synthetically generate more diverse unbound structures using physical simulations or deep learning tools. Similar to previous work [27], we define the interpolation process in ℝ^3^ and *SO*(3) after a global RMSD alignment anchored on the receptor protein. The conditional probability paths for translation, rotation, and internal flexibility are thus:

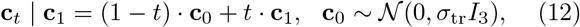

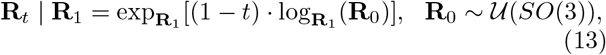

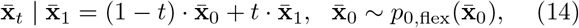

where **c** is the center of mass of the ligand protein, **R** is the rotation matrix of the ligand protein with regard to a reference frame, and 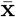 is the all-atom coordinates of the receptor and ligand proteins after individual RMSD alignment. Although we interpolate all-atom coordinates, AF2Dock only utilizes the backbone and C_*β*_ atoms during embedding and thus only learns a distribution over those atoms. The pseudocode for the training and inference of AF2Dock is in the Supplemental Material, Algorithms S5 and S7.

### C. Data

We train AF2Dock on the PINDER training set [21], an annotated dataset of 2.3 million bi-protein complexes that uses structural clustering to derive interface-based data splits. In addition to holo monomer structures (conformation in the bound experimental complex structure), PINDER also includes apo (experimentally resolved unbound protein conformation) and AF2-predicted monomer structures of a protein chain when available, allowing learning of flexible docking. In addition to bi-protein complexes in the PINDER dataset, we also construct tri-protein complexes for our training set. Additional details on training set filtering and tri-protein complex construction are in the Supplemental Material.

We use two test sets to evaluate model performance. The first is the PINDER-AF2 benchmark [21], which was filtered to have no structural interface cluster overlap with the training sets of AF-M and AF3. This test set includes 180 complexes with holo monomer structures, 30 complexes with apo monomer structures, and 127 complexes with predicted monomer structures. The second test set is an antibody nanobody set [22] that was previously used to evaluate the performance of AF3, which was filtered against AF-M’s and AF3’s training set using release dates and sequence identities of the CDR loops. This set contains 49 antibody complexes and 60 nanobody complexes. In addition to the holo input structures, we also construct predicted input structures for this set by running AF3 on antibody/nanobody and antigen sequences individually with one seed and using the top-ranked prediction. We only perform two-body docking tests in this work. Therefore, for antibodies, we combine the heavy and light chains into a single chain with an added index gap in between chains and dock it as a single entity (Supplemental Material).

### D. Model training and inference

We implement AF2Dock based on the OpenFold implementation [20] of AF-M in PyTorch. We train the model in three stages. During stage one, we only use holo monomer structure inputs and train only the docking module while freezing the weights of AF-M components. The reason for using only holo monomers in this stage is due to the training instability that we observed. For stage two, we fine-tune the docking module while still keeping the weights of AF-M components frozen. In this stage, for each protein chain, we randomly sample holo, apo, and predicted monomer structures with a probability of 20%, 40%, and 40%, when available. Finally, for stage three, we perform full-parameter fine-tuning of AF-M along with the docking module, while using the same monomer sampling scheme as stage two.

During inference of AF2Dock, for each sample, we integrate over 10 time steps along the learned velocity field by default. We sample 20 structures for the PINDER-AF2 benchmark and 40 structures for the antibody/nanobody test set. We rank the resulting predictions by their ipTM score. We compare AF2Dock with various baseline methods, including single-sequence AF-M with custom template inputs [1], co-folding models including standard AF-M [1] and AF3 [2], diffusion-based docking models DiffDock-PP [12] and DFMDock [13], as well as physics-based docking methods HDock [28] and ZDock [29]. For co-folding AF-M, AF3, DiffDock-PP, and DFMDock, we sample the same number of structures as AF2Dock. Additional details regarding training and inference are available in the Supplemental Material.

## III. RESULTS

### A. AF2Dock performs strongly against baseline structure-based docking methods while producing orthogonal predictions to co-folding models on the PINDER-AF2 benchmark

To evaluate the performance of AF2Dock and compare it to various baseline methods, we tested them on the PINDER-AF2 benchmark, which was constructed to have no structural interface overlap with the training sets of AF-M and AF3 and includes holo (conformation in the bound experimental structure), apo (experimentally resolved unbound monomer structures), and AF2-predicted monomer structures for many test targets. We compare AF2Dock with both structure-based docking methods and co-folding methods, including single-sequence AF-M with custom template inputs (referred to as single-sequence AF-M herein), standard co-folding AF-M and AF3, diffusion-based docking models DiffDock-PP and DFMDock, and physics-based docking methods HDock and ZDock. For co-folding AF-M and AF3, we use the standard template pipeline and do not input the same templates as the structure-based docking methods. We compare the prediction accuracy by computing the success rates of the Top-1 and Top-5 ranked predictions, as well as the oracle success rate that corresponds to the best among all predicted structures regardless of their ranking (Fig. 2).

**FIG. 2.**
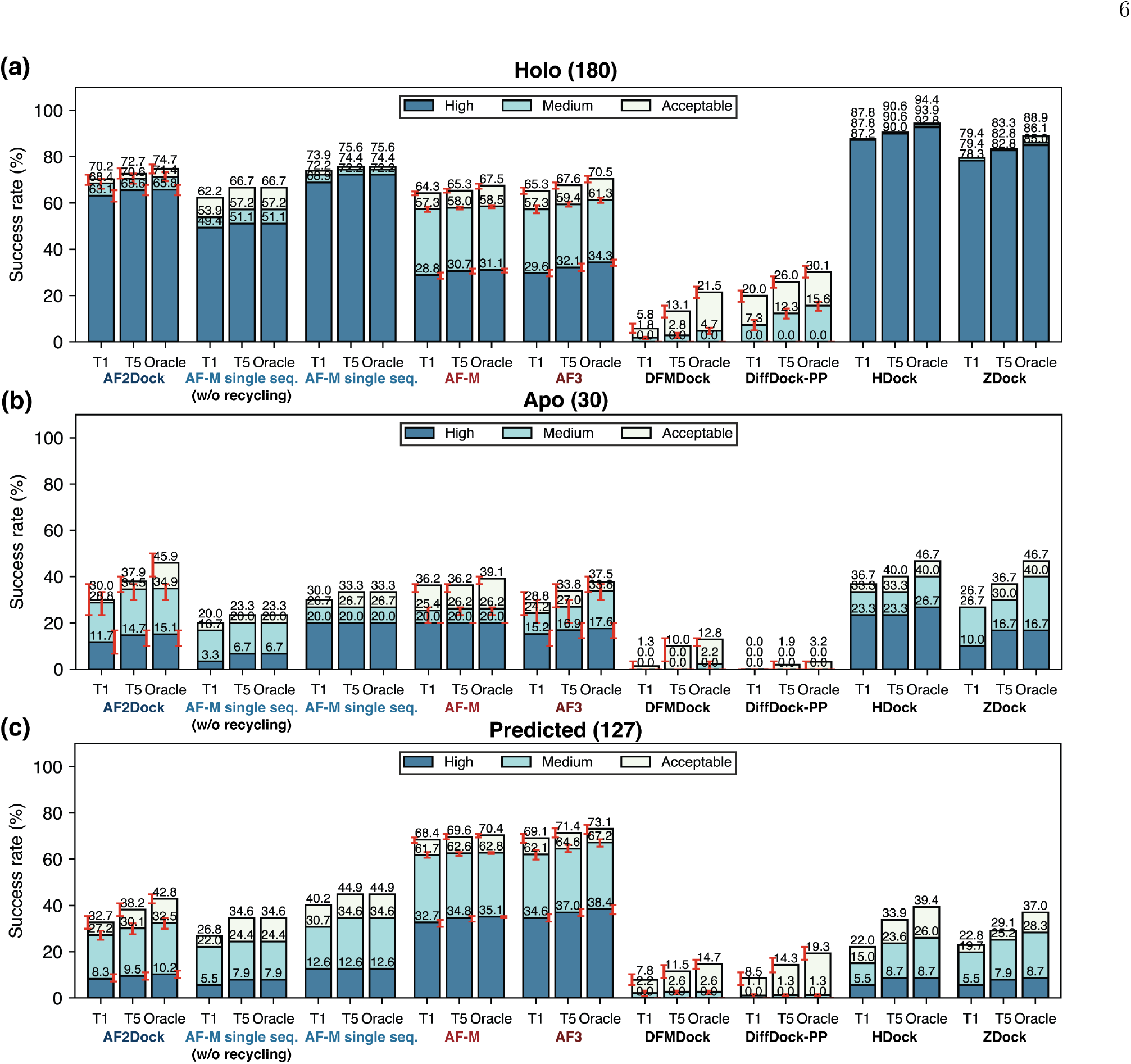
Success rates on the PINDER-AF2 test set. Shown are Top-1 (T1), Top-5 (T5), and oracle success rates of AF2Dock, single-sequence AF-M, co-folding AF-M, AF3, DFMDock, DiffDock-PP, HDock, and ZDock on the PINDER-AF2 benchmark when using (a) holo, (b) apo, and (c) predicted monomer structures in docking. The oracle category includes five predictions for single-sequence AF-M and 20 predictions for other methods. Bar heights and error bars represent the mean and the 95% confidence interval of 10,000 bootstrap samples. We do not compute uncertainties for single-sequence AF-M, HDock, and ZDock, as there is no stochasticity in their sampling process. Docking quality is defined by DockQ thresholds, with acceptable *>* 0.23, medium *>* 0.49, and high *>* 0.80.

When supplied with holo monomer structures as templates, which are the exact conformation that the sub-units adopt in the bound complex, single-sequence AF-M (when using default recycling iterations) surprisingly outperforms co-folding AF-M, showing a large percentage of high-quality predictions (Fig. 2a). This strong performance suggests that, although trained with MSA inputs, AF-M has learned to predict inter-protein contacts based solely on structural features. This result may also explain AF-M’s relatively high performance when using unpaired MSAs, where AF-M likely infers inter-chain contacts based on per-chain structural features derived from intra-chain MSAs, instead of relying on inter-chain co-evolution. When we perform predictions with AF2Dock using holo structure inputs, we obtain success rates higher than those of single-sequence AF-M without using recycling iterations (as is the case for AF2Dock), but slightly lower than those of single-sequence AF-M with recycling (Fig. 2a). This result suggests that the recycling-based refinement still performs better than our flow-matching-based scheme in the case of holo inputs. When we compare the accuracy of AF2Dock and single-sequence AF-M with other structure-based docking methods, we observe that while they still fall short of physics-based docking methods, they substantially outperform existing diffusion-based docking models, greatly bridging the performance gap between deep-learning-based and physics-based docking methods in the holo setting.

Although the strong performance of AF2Dock and single-sequence AF-M using holo inputs is encouraging, holo structures are unavailable for unsolved targets in real-world settings. As a result, we evaluate model performance with apo and predicted structure inputs that in principle can be obtained without access to the bound structure. The results (Fig. 2b-c) show that although AF2Dock can still predict correct structures for certain targets, the success rates drop substantially compared to using holo structures. The overall lower success rates for the apo test set compared to holo and predicted sets among all methods, including co-folding models, may result from the low number of targets in this set (30), which can happen to be skewed towards targets with more dissimilar interfacial features compared to the training set. The trend of decreasing success rate using non-holo inputs is shared among all structure-based docking methods, with AF2Dock performing competitively or outperforming all other docking methods in this setting. In contrast, AF2Dock underperforms co-folding AF-M and AF3, especially when using predicted monomer structures as inputs. This is likely caused by the conformational changes required at the binding interface for successful docking and suggests that AF2Dock cannot effectively explore the flexibility of the input monomer structures.

To examine the ability of AF2Dock to model internal flexibility, we compare the per-subunit interface RMSD (ps-iRMSD, computed for interface residues in each subunit separately with each subunit individually aligned) among input, output, and ground truth (GT) structures when using predicted monomer inputs (Fig. 3a). The input−output ps-iRMSD is much smaller compared to input−GT ps-iRMSD, suggesting that the internal structural movements induced by AF2Dock is minimal compared to the required changes to reach the GT structure. In addition, the difference between input − GT and output − GT ps-iRMSD is not statistically significant based on a one-sided Wilcoxon signed-rank test, indicating that AF2Dock does not consistently produce internal structural movements towards the correct bound conformation. When we separate the ps-iRMSD of successful predictions based on their difficulty category (as defined in Docking Benchmark 4.0 [30]), we observe a larger difference between input − GT and output −GT ps-iRMSD with higher difficulty (Fig. S1), although still none of them is statistically significant. This inability to properly model subunit flexibility is further reflected in the success rates of each difficulty category (Fig. 3b-c), where AF2Dock shows a steep decrease in prediction success rates going from rigid-body to difficult targets, while co-folding AF-M only drops moderately, especially for acceptable-quality predictions.

**FIG. 3.**
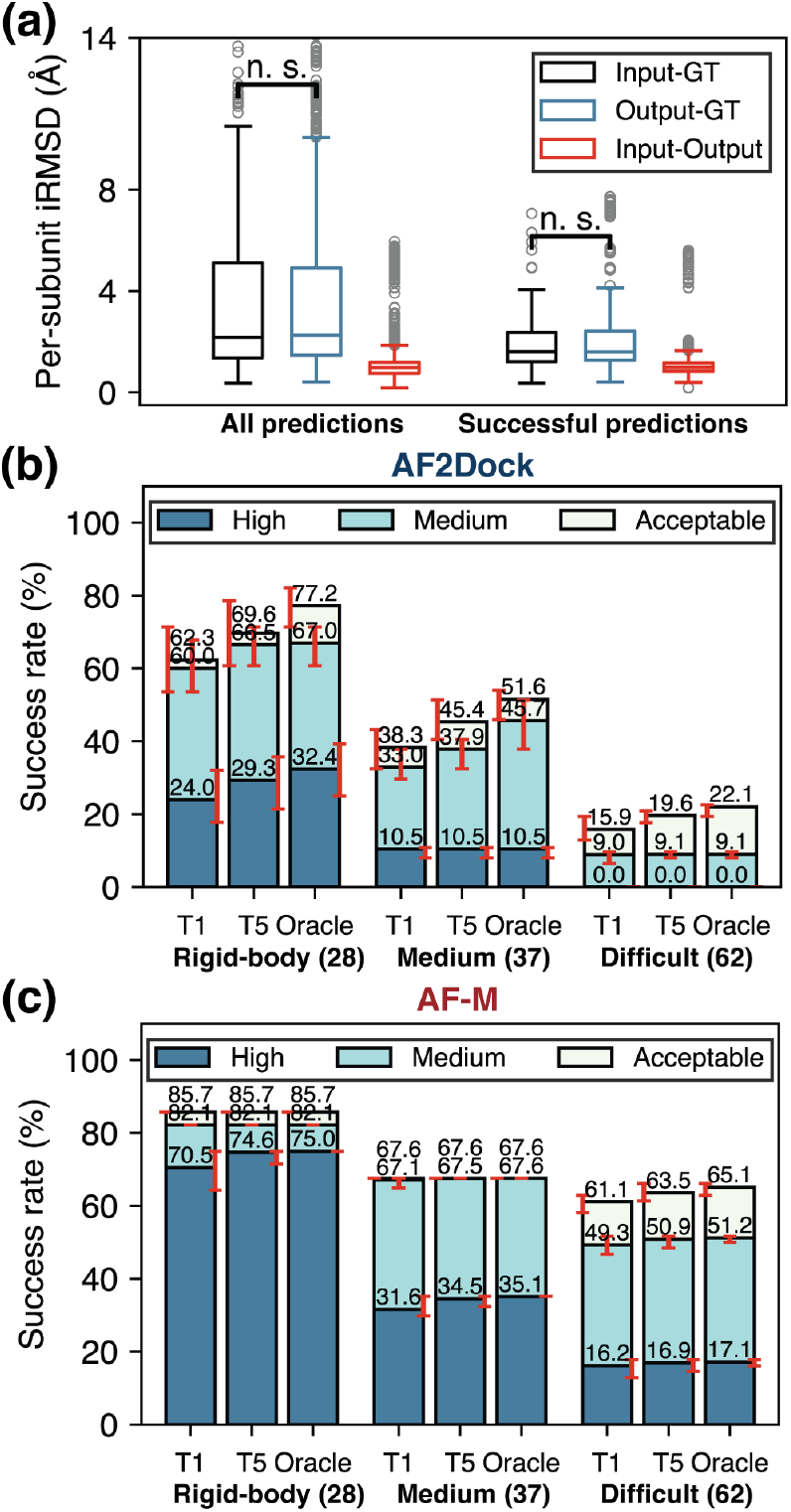
AF2Dock cannot effectively explore subunit flexibility. (a) Per-subunit interface RMSD (ps-iRMSD) among input, output, and ground truth (GT) structures for all predictions and successful predictions of AF2Dock on the PINDER-AF2 predicted set. Successful predictions are ones with DockQ scores *>* 0.23. One-sided Wilcoxon signed-rank test was performed between input-GT and output-GT ps-iRMSD. (b-c) Top-1 (T1), Top-5 (T5), and oracle success rates of (b) AF2Dock and (c) co-folding AF-M on different difficulty categories of the PINDER-AF2 benchmark using predicted inputs. The oracle category includes 20 predictions for both methods. Bar heights and error bars represent the mean and the 95% confidence interval of 10,000 bootstrap samples. Docking quality is defined by DockQ thresholds, with acceptable *>* 0.23, medium *>* 0.49, and high *>* 0.80.

When we plot the highest DockQ scores from the Top-1, Top-5, and oracle categories of predictions for each target by AF2Dock and co-folding AF-M against each other, we observe that the predictions of the two methods are orthogonal (Fig. 4), as evidenced by the presence of targets in the top-left and bottom-right quadrants. Although we also observe similar orthogonality between single-sequence AF-M and co-folding AF-M using holo inputs (Fig. S2a), the number of orthogonal predictions of single-sequence AF-M is lower than that of AF2Dock when using apo or predicted inputs (Fig. S2b-c), despite single-sequence AF-M showing a comparable or higher success rate in certain cases. This higher orthogonality of AF2Dock compared to single-sequence AF-M suggests that AF2Dock may be better suited to complement co-folding methods by combining predictions from different methods, which may lead to higher overall success rates. However, a naive combination of AF2Dock and co-folding AF-M predictions followed by re-ranking using the ipTM score only leads to improved Top-1 and Top-5 success rates in the case of holo inputs (Fig. S3), suggesting over-confident ipTM scores from both methods. More sophisticated combination schemes or manual inspection of the predicted structures may be required to harness this orthogonality between AF2Dock and co-folding AF-M.

**FIG. 4.**
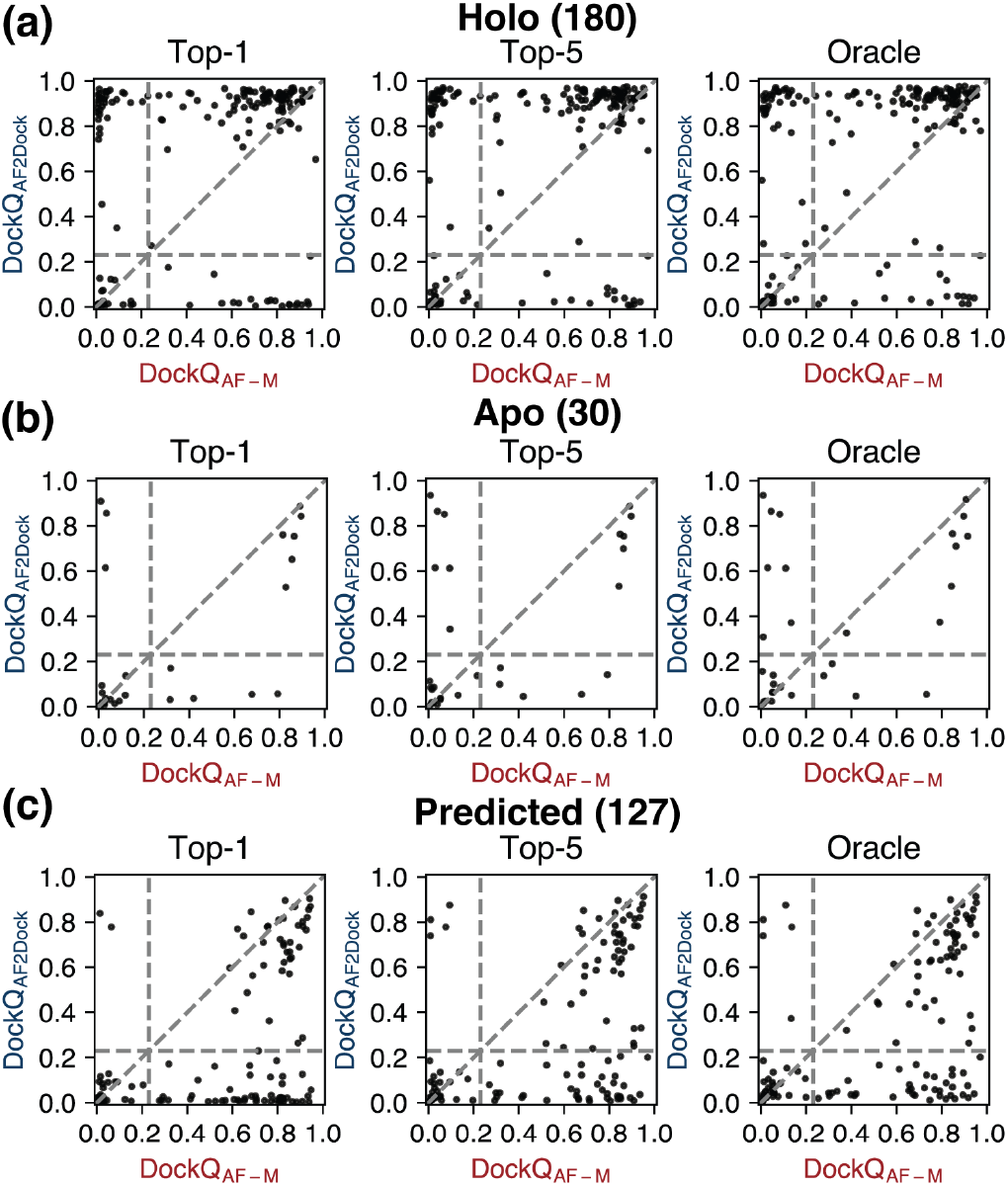
Orthogonality in successful predictions between AF2Dock and co-folding AF-M. Highest DockQ scores from the Top-1, Top-5, and oracle categories of predictions by AF2Dock and co-folding AF-M plotted against each other for the PINDER-AF2 benchmark when using (a) holo, (b) apo, and (c) predicted structures in docking. Vertical and horizontal lines at DockQ = 0.23 (acceptable docking quality threshold), as well as the line DockQ_AF2Dock_ = DockQ_AF-M_ are shown in gray as visual guides.

### B. AF2Dock successfully predicts antibody and nanobody complexes but shows poor ranking when using predicted inputs

In contrast to other naturally evolved protein-protein interactions, antibodies are selected against antigens in the immune system through a process that rarely involves the co-evolution of the antigens. In addition, different antibodies may target different epitopes of an antigen, while other natural protein-protein interactions are typically conserved in locality throughout evolution. As a result, MSAs of antibodies and antigens contain limited, if any, co-evolutionary signals regarding inter-chain contacts, resulting in poor performance of co-folding models on these targets. To test whether using structure-based features instead of MSAs could benefit these challenging protein complexes, we performed predictions with AF2Dock as well as baseline co-folding and docking methods on an antibody/nanobody test set that was previously used to benchmark the performance of AF3 [22] (Fig. 5).

**FIG. 5.**
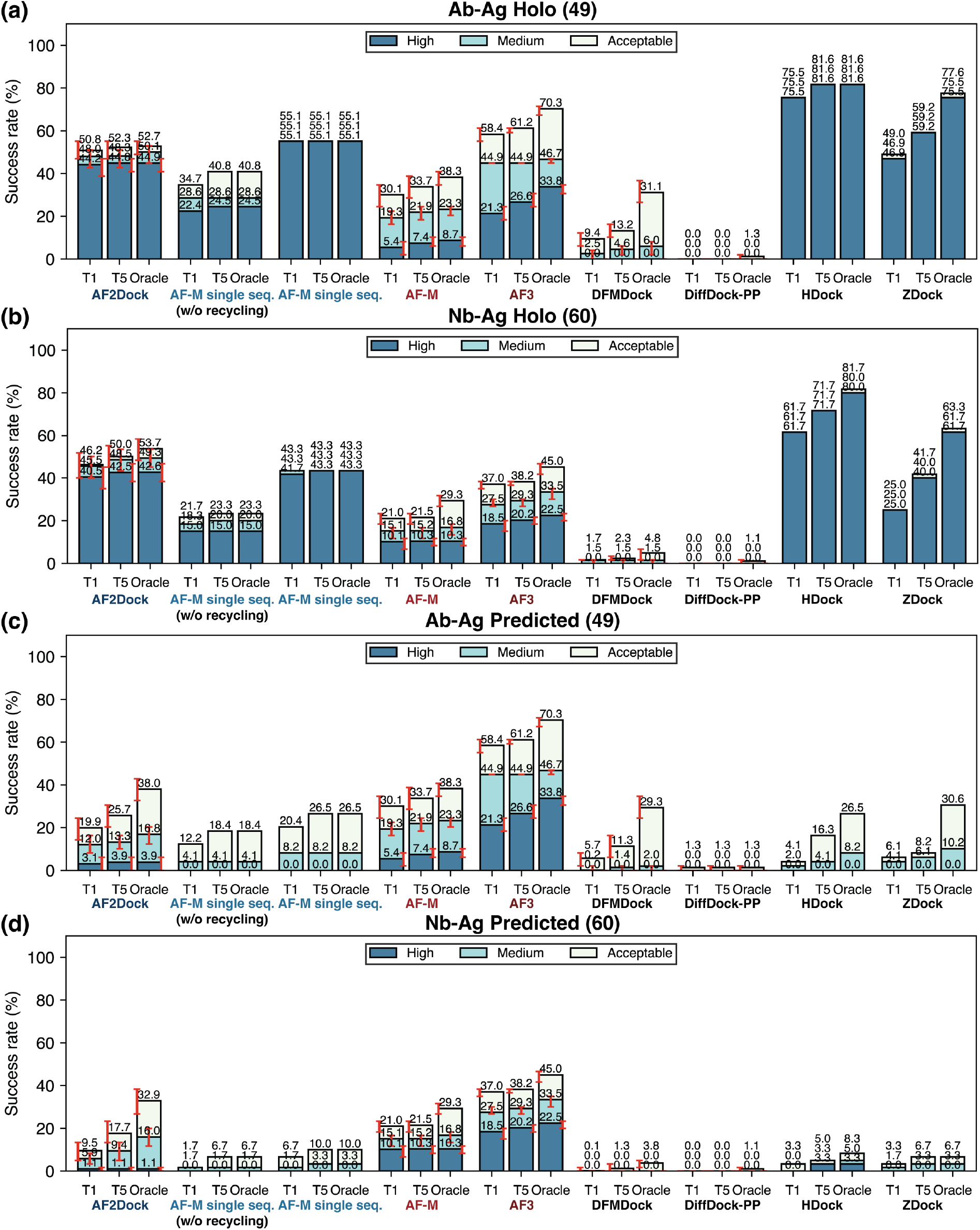
Success rates on the antibody/nanobody test set. Shown are Top-1 (T1), Top-5 (T5), and oracle success rates of AF2Dock, single-sequence AF-M, co-folding AF-M, AF3, DFMDock, DiffDock-PP, HDock, and ZDock on (a,c) antibody-antigen (Ab-Ag) complexes as well as (b,d) nanobody-antigen (Nb-Ag) complexes with (a,b) holo and (c,d) predicted inputs in docking. The oracle category includes five predictions for single-sequence AF-M and 40 predictions for other methods. Bar heights and error bars represent the mean and the 95% confidence interval of 10,000 bootstrap samples. We do not compute uncertainties for single-sequence AF-M, HDock, and ZDock, as there is no stochasticity in their sampling process. Docking quality is defined by DockQ thresholds, with acceptable *>* 0.23, medium *>* 0.49, and high *>* 0.80.

The results reveal a similar trend as the PINDER-AF2 benchmark, where AF2Dock performs substantially better with holo input structures compared to using predicted inputs (Fig. 5). Notably, when using predicted inputs, AF2Dock outperforms all other structure-based docking methods tested here on nanobody complexes, including single-sequence AF-M and physics-based docking methods (Fig. 5c-d). Comparing with co-folding models, however, AF2Dock still underperforms both AF-M and AF3 (Fig. 5c-d). Interestingly, we observe that the oracle success rates of AF2Dock using predicted inputs is similar to that of co-folding AF-M, yet its Top-1 and Top-5 success rates are lower, implicating a failure to accurately rank the predictions, again suggesting that the ipTM score is overconfident in false positive structures. Additionally, there is a larger performance gap between AF2Dock and physics-based docking methods on the antibody/nanobody test set compared to the PINDER-AF2 set when using predicted inputs. Although this difference may be reflective of improved effectiveness of AF2Dock compared to physics-based methods specifically on loop-based interactions, we note that it could also be potentially due to data leakage, as the antibody/nanobody test set was filtered using less stringent criteria [22] compared to the PINDER-AF2 benchmark [21], mainly because antibody sequences and structures are highly homologous while data are scarce.

When we compare the highest DockQ score among the Top-1, Top-5, and oracle categories of predictions between methods, we again observe orthogonality between AF2Dock and AF-M (Fig. S4), as well as AF3 (Fig. S5, Top-1 with predicted inputs shown in Fig. 6), even when using predicted input structures. Similar to the PINDER-AF2 benchmark, although single-sequence AF-M also exhibits some orthogonality with co-folding models, the number of orthogonal predictions is smaller than that of AF2Dock (Fig. S6-S7). When we inspect cases where AF2Dock was successful while AF3 failed (Fig. 6), we observe that the AF3 predicted structures are often localized in a small region, likely resulting from spurious signals from the input MSAs or biases in the training data. In contrast, AF2Dock is able to explore a wider variety of binding locations resulting in successful predictions. Nevertheless, a naive combination of predictions between AF2Dock and AF-M followed by re-ranking still did not improve the Top-1 and Top-5 success rates (Fig. S8).

**FIG. 6.**
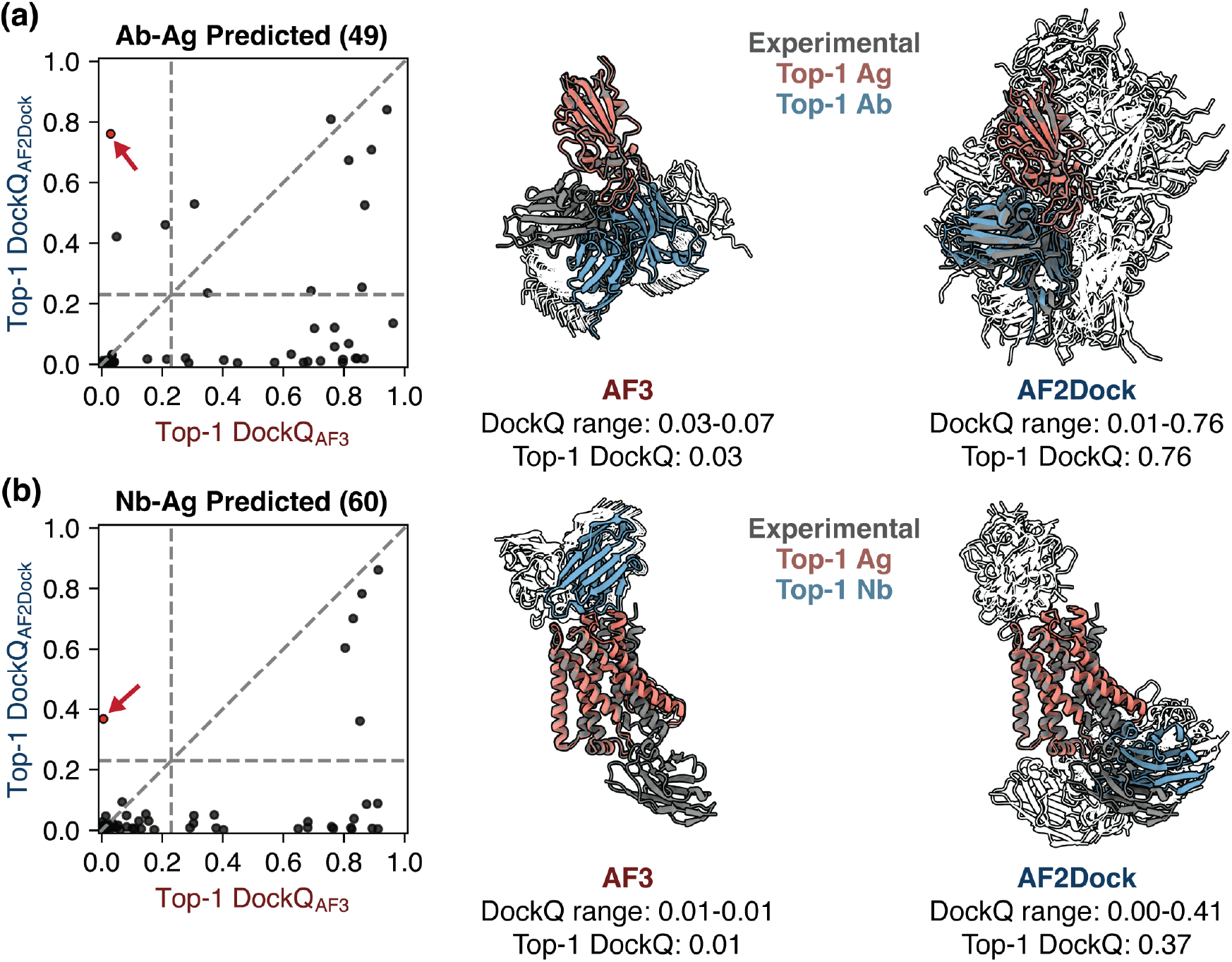
AF2Dock produces orthogonal predictions compared to AF3 on antibody and nanobody complexes even when using predicted inputs. Left panels show DockQ scores of the Top-1 predictions from AF2Dock using predicted inputs and AF3 plotted against each other for (a) antibody-antigen (Ab-Ag) and (b) nanobody-antigen (Nb-Ag) complexes. Vertical and horizontal lines at DockQ = 0.23 (acceptable docking quality threshold), as well as the line DockQ_AF2Dock_ = DockQ_AF-M_ are shown in grey as visual guides. On the right, example orthogonal predictions between AF3 and AF2Dock (red data points in the left panels) are shown for (a) an antibody-antigen complex (PDB: 8qrg) and (b) a nanobody-antigen complex (PDB: 7zk1). Experimental ground truth structures are shown as gray, and Top-1 predictions of each method are shown as salmon (Ag) and blue (Ab/Nb). All lower-ranked predictions are shown as silhouettes in the background.

We also performed preliminary tests on whether applying external scores on AF2Dock predictions could improve ranking (Supplemental Material). We tested the three scoring methods: Rosetta interface score *I*_*sc*_ [31], Piston score [32], and AF-M ipTM with template input in single-sequence mode (which we dub AFMRank, similar to AF2Rank [16]). While the scores tested here all exhibit the ability to distinguish good versus bad structures, they all produce false positives when using a static score cutoff (Fig. S15). When applied to ranking, we find that although individual scoring metrics did not result in substantially different success rates compared to the default AF2Dock ranking (Fig. S17), they succeeded in ranking different subsets of targets (Fig. S18 - S21). Thus, a future direction is to explore the combination of multiple ranking metrics to allow improved ranking.

When inspecting the predicted input structures of antibodies, we find that in many cases the complementary determining region 3 of the heavy chain (CDRH3), typically involved in antigen interactions, is in a very different conformation compared to the bound structure (Fig. S9a). This structural error leads to clashes when the two subunits are in the correct relative orientation (Fig. S9a), which is a likely cause of failure in docking predictions because AF2Dock cannot effectively explore subunit flexibility as shown in the previous section. One advantage of AF2Dock compared to other structure-based docking methods, owing to its adaptation from the AF-M architecture, is the ability to inpaint missing residues. Therefore, we tested docking using input structures with the CDRH3 loop and low pLDDT regions removed, such that a more correct conformation may be inpainted with a proper structural context. When we compare the DockQ scores of the best predictions for each target between using full structural inputs and inpainting (Fig. S10a-b), we find that the effect of the inpainting strategy is inconsistent, showing improvement in some targets while hurting performance in others. In terms of success rates, for antibody complexes (Fig. S10c, left panel), although the Top-1 acceptable success rate stays the same when using the inpainting strategy, the medium and Top-5 success rates improve. When we combine predictions from the two strategies and and re-rank with ipTM, we obtain further improved overall success rates compared to only using full structural inputs (Fig. S10c, left panel). When we inspect an antibody-antigen target where the inpainting strategy leads to better predicted structures (Fig. S9c), we find that removing the CDRH3 loop from the input indeed significantly increased the conformational diversity of that region in the output structures (Fig. S9b). Interestingly, when the inpainting strategy is applied to nanobody complexes, we observe a substantial decrease in success rates (Fig. S10c, right panel), although the combination of predictions between strategies still leads to a slightly improved Top-1 success rate. This contrast in performance compared to the antibody cases is perhaps because CDRH3 is often solely responsible for mediating interactions with antigens in nanobodies, such that omitting it from the input structure causes AF2Dock to fail to identify the correct binding interface.

### C. Ablation studies reveal the importance of full-parameter fine-tuning and nuances in the role of input features

To understand the impact of incorporating ESM embeddings in the docking module as well as full-parameter fine-tuning in stage three of training, we performed ablation studies where we omitted these elements and tested the performance of the resulting models on both test sets described in the previous sections (Fig. S11-S12)

When we omit full-parameter fine-tuning (Fig. S11-S12), although the model still outperforms single-sequence AF-M without recycling, it substantially underperforms base AF2Dock, particularly in Top-1 performance. This result suggests that by fine-tuning AF-M components along with the docking module, we allow information to better flow throughout the model, leading to improved structure prediction and confidence ranking.

When we train a model variant without using ESM embeddings of individual proteins in the docking module, we surprisingly find that the non-ESM variant of AF2Dock performs better compared to base AF2Dock when using holo structure inputs (Fig. S11a and Fig. S12a-b). This result suggests that the features present in the ESM embeddings may mislead predictions for certain complexes compared to solely using structure-based features, as the model may learn to rely on spurious signals in ESM embeddings that are not generalizable to new targets. Additionally, since we only train the docking module from scratch while fine-tuning the AF-M components, which do not incorporate ESM embeddings natively, our model may not be able to fully utilize the information in the embeddings. In contrast to the holo set, however, we find that the non-ESM variant of AF2Dock performs similarly to base AF2Dock on the apo and predicted PINDER-AF2 test sets (Fig. S11b-c). In the case of antibody/nanobody complexes using predicted inputs, we find an interesting pattern where base AF2Dock performs better on the antibody complexes, while the non-ESM variant performs better (in Top-1 and Top-5 success rates) on the nanobody complexes (Fig. S12c-d, nanobody success rate comparison also shown in Fig. 7a). This discrepancy between antibodies and nanobodies may be due to the difference in their interface sizes. With a larger interface (antibodies), the sequence features from ESM embeddings could potentially be helpful for the model to identify epitope/paratopes, while with a small interface (nanobodies), sequence features may be misleading, and structure information alone may be more indicative. The discrepancy may also be reflective of the fact that there are more antibody training examples than nanobody ones, which could cause the model to be more likely to overfit on spurious signals in the ESM embeddings for nanobodies. When we compare the highest DockQ scores among the Top-1, Top-5, and oracle categories between predictions by the non-ESM variant of AF2Dock and AF3 (Fig. S13, Top-1 comparison for nanobody using predicted inputs shown in Fig. 7b), we still observe orthogonality in prediction success and a higher number of orthogonally successful predictions (top left quadrant) compared to base AF2Dock (Fig. 6b). These trends suggest nuances in incorporating features for structure-based docking models, where more features may not lead to improved performance in all cases, and a model ensembling approach may provide a more robust outcome.

**FIG. 7.**
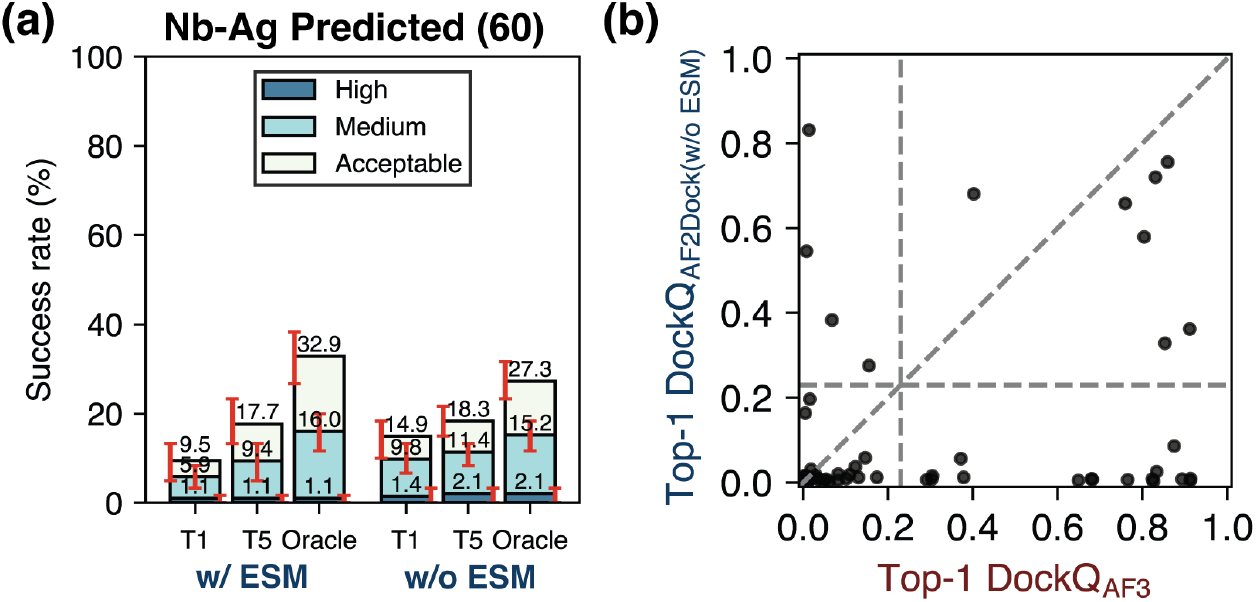
The non-ESM variant of AF2Dock performs better than base AF2Dock for nanobody-antigen (Nb-Ag) complexes. (a) Top-1 (T1), Top-5 (T5), and oracle success rates of base AF2Dock (with ESM) and the non-ESM variant on the nanobody test set using predicted inputs. The oracle category includes 40 predictions. Bar heights and error bars represent the mean and the 95% confidence interval of 10,000 bootstrap samples. Docking quality is defined by DockQ thresholds, with acceptable *>* 0.23, medium *>* 0.49, and high *>* 0.80. (b) DockQ scores of the Top-1 predictions from the non-ESM variant of AF2Dock using predicted inputs and AF3 plotted against each other on the nanobody test set. Vertical and horizontal lines at DockQ = 0.23 (acceptable docking quality threshold), as well as the line DockQ_AF2Dock_ = DockQ_AF-M_ are shown in grey as visual guides.

Finally, we explore how the use of full-complex templates and the interpolation process may have contributed to the success of AF2Dock compared to single-sequence AF-M. We performed inference of two variants of single-sequence AF-M without recycling on the PINDER-AF2 benchmark, with one variant using noisy full-complex templates as inputs, and the other using the same interpolation process as AF2Dock on top of using full-complex templates (Figure S14). We find that although the interpolation process further improved the success rates of non-recycling single-sequence AF-M even when using vanilla AF-M architecture and weights, the performance is still substantially inferior compared to AF2Dock and single-sequence AF-M with recycling, suggesting merits in our docking module and flow-matching-based training and inference scheme.

## IV. DISCUSSION

In this work, we present a novel method for adapting co-folding models for structure-based protein-protein docking through a novel docking module and a flow-matching-based training and inference scheme. Applying our methodology to AF-M, we show that the resulting model AF2Dock, when using non-holo structure inputs, performs competitively with baseline structure-based docking methods and, in the case of nanobody complexes, outperforms all other docking methods tested here. We also provide, for the first time, a comprehensive evaluation of single-sequence AF-M for structure-based protein-protein docking, which reveals its surprisingly strong performance, especially in the holo setting. When compared to co-folding models, AF2Dock still underperforms both AF-M and AF3 when using predicted inputs, although we show the presence of orthogonally successful predictions between the two methods. The orthogonality suggests the possibility of improved results by combining AF2Dock with co-folding methods, yet a naive combination of AF2Dock’s and AF-M’s predictions followed by re-ranking with ipTM did not result in improved Top-1 and Top-5 success rates in most cases, suggesting improper ranking with the confidence score.

A key limitation of AF2Dock is the inability to properly explore subunit flexibility, which limits its performance on targets where binding involves substantial conformational changes. Conformational change during binding has been recognized as the limiting factor in protein-protein docking for a long time [33, 34]. Here, although we attempt to explicitly model internal flexibility during training and inference, the performance of AF2Dock on this aspect is unsatisfactory. This limitation could result from various reasons, including the limited variability in training input structures, as well as the intrinsic tendency of AF-M to retain template inputs in single-sequence mode. Future directions for improving flexibility modeling include training with more diverse input structures obtained from physical simulations or deep learning predictions, or using a more well-defined internal flexibility prior. In addition, using an AF3-like architecture instead of AF-M may benefit the exploration of conformational diversity, as the diffusion-based structural generation in AF3 is less deterministic compared to the regression-based AF-M.

Although in this work we mainly explore structure-based docking without MSAs, MSAs can provide valuable information in many cases regarding subunit flexibility [35, 36] and inter-chain contacts. A naive inclusion of MSAs features in AF2Dock did not lead to improved flexibility or performance (not shown). A future direction is to explore how to better combine both structure-based features and MSAs while avoiding their shortcomings to improve upon the state-of-the-art AF3. One possibility is to devise an architecture where MSAs are only used for intra-chain folding while using structure-based features for predicting inter-chain contacts, which would allow better exploration of subunit flexibility while being unaffected by spurious inter-chain signals in MSAs.

Finally, accurate ranking of the predicted structures continues to be a challenging task for docking methods with the sample-and-rank scheme. From our results, although the ipTM score correlates well with prediction quality, it is not perfect and can be overconfident in false positive predictions, hindering the combination of results from AF2Dock and AF-M. Additionally, our preliminary tests on external scoring methods suggest that individual scores do not provide substantially different ranking performance compared to AF2Dock ipTM, yet different scores successfully rank different subsets of the targets. A future direction is thus to devise a better method for ranking predictions, either by improving model scoring output through using alternative scores like ipSAE [37], or by exploring the combination of multiple ranking metrics. In the case of AF2Dock, another source for ranking difficulty comes from incorrect interface-residue conformations even with correct relative subunit orientations. Strategies similar to AlphaRED [38] where physics-based local relaxation is performed after deep-learning-based predictions may help to better model induced-fit, which can lead to improved predictions and ranking.

## V. ACKNOWLEDGMENTS

This work was supported by National Institutes of Health grant R35-GM141881 and by AstraZeneca. Computational resources were provided by the Advanced Research Computing at Hopkins (ARCH).

## VI. DATA AVAILABILITY

Code for AF2Dock is available on GitHub (https://github.com/Graylab/AF2Dock). Model weights are available in Zenodo entry 17782958 [39]. Training and test sets are available from prior publications [21, 22]. AF3-predicted input structures for the antibody/nanobody test set are available in Zenodo entry 17782958 [39].

## References

[1] R. Evans, M. O’Neill, A. Pritzel, N. Antropova, A. Senior, T. Green, A. Žídek, R. Bates, S. Blackwell, J. Yim, O. Ronneberger, S. Bodenstein, M. Zielinski, A. Bridgland, A. Potapenko, A. Cowie, K. Tunyasuvunakool, R. Jain, E. Clancy, P. Kohli, J. Jumper, and D. Hassabis, Protein complex prediction with AlphaFold-multimer, bioRxiv, 2021.10.04.463034 (2021).

[2] J. Abramson, J. Adler, J. Dunger, R. Evans, T. Green, A. Pritzel, O. Ronneberger, L. Willmore, A. J. Ballard, J. Bambrick, S. W. Bodenstein, D. A. Evans, C.-C. Hung, M. O’Neill, D. Reiman, K. Tunyasuvunakool, Z. Wu, A. Žemgulytė, E. Arvaniti, C. Beattie, O. Bertolli, A. Bridgland, A. Cherepanov, M. Congreve, A. I. Cowen-Rivers, A. Cowie, M. Figurnov, F. B. Fuchs, H. Gladman, R. Jain, Y. A. Khan, C. M. R. Low, K. Perlin, A. Potapenko, P. Savy, S. Singh, A. Stecula, A. Thillaisundaram, C. Tong, S. Yakneen, E. D. Zhong, M. Zielinski, A. Žídek, V. Bapst, P. Kohli, M. Jaderberg, D. Hassabis, and J. M. Jumper, Accurate structure prediction of biomolecular interactions with AlphaFold 3, Nature 630, 493 (2024).

[3] C. Discovery, J. Boitreaud, J. Dent, M. McPartlon, J. Meier, V. Reis, A. Rogozhnikov, and K. Wu, Chai-1: Decoding the molecular interactions of life, bioRxiv (2024).

[4] S. Passaro, G. Corso, J. Wohlwend, M. Reveiz, S. Thaler, R. Somnath, N. Getz, T. Portnoi, J. Roy, H. Stark, D. Kwabi-Addo, D. Beaini, T. Jaakkola, and R. Barzilay, Boltz-2: Towards accurate and efficient binding affinity prediction, bioRxiv (2025).

[5] X. Chen, Y. Zhang, C. Lu, W. Ma, J. Guan, C. Gong, J. Yang, H. Zhang, K. Zhang, S. Wu, K. Zhou, Y. Yang, Z. Liu, L. Wang, B. Shi, S. Shi, and W. Xiao, Protenix - advancing structure prediction through a comprehensive AlphaFold3 reproduction, bioRxiv (2025).

[6] N. Corley, S. Mathis, R. Krishna, M. S. Bauer, T. R. Thompson, W. Ahern, M. W. Kazman, R. I. Brent, K. Didi, A. Kubaney, L. McHugh, A. Nagle, A. Favor, M. Kshirsagar, P. Sturmfels, Y. Li, J. Butcher, B. Qiang, L. L. Schaaf, R. Mitra, K. Campbell, O. Zhang, R. Weissman, I. R. Humphreys, Q. Cong, J. Funk, S. Sonthalia, P. Liò, D. Baker, and F. DiMaio, Accelerating biomolecular modeling with AtomWorks and RF3, bioRxiv (2025).

[7] M. L. Rennie and M. R. Oliver, Emerging frontiers in protein structure prediction following the AlphaFold revolution, J. R. Soc. Interface 22, 20240886 (2025).

[8] F. Costa, M. Blum, and A. Bateman, Keeping it in the family: using protein family templates to rescue low confidence AlphaFold2 models, Bioinform. Adv. 4, vbae188 (2024).

[9] R. Yin, B. Y. Feng, A. Varshney, and B. G. Pierce, Benchmarking AlphaFold for protein complex modeling reveals accuracy determinants, Protein Sci. 31, e4379 (2022).

[10] Z. Zhang, H. K. Wayment-Steele, G. Brixi, H. Wang, D. Kern, and S. Ovchinnikov, Protein language models learn evolutionary statistics of interacting sequence motifs, Proc. Natl. Acad. Sci. U. S. A. 121, e2406285121 (2024).

[11] I. A. Vakser, Protein-protein docking: From interaction to interactome, Biophys. J. 107, 1785 (2014).

[12] M. A. Ketata, C. Laue, R. Mammadov, H. Stärk, M. Wu, G. Corso, C. Marquet, R. Barzilay, and T. S. Jaakkola, DiffDock-PP: Rigid protein-protein docking with diffusion models, arXiv [q-bio.BM] (2023).

[13] L.-S. Chu, S. Sarma, and J. J. Gray, Unified sampling and ranking for protein docking with DFMDock, bioRxiv (2024).

[14] M. McPartlon, C. Marquet, T. Geffner, D. Kovtun, A. Goncearenco, Z. Carpenter, L. Naef, M. M. Bronstein, and J. Xu, Bridging sequence and structure: Latent diffusion for conditional protein generation, Machine Learning in Structural Biology Workshop (2023).

[15] F. Sverrisson, M. Akdel, D. Abramson, J. Feydy, A. Goncearenco, Y. Adeshina, D. Kovtun, C. Marquet, X. Zhang, D. Baugher, Z. Carpenter, L. Naef, M. M. Bronstein, and B. Correia, DiffMaSIF: Score-based diffusion models for protein surfaces, Machine Learning in Structural Biology Workshop (2023).

[16] J. P. Roney and S. Ovchinnikov, State-of-the-art estimation of protein model accuracy using AlphaFold, Phys. Rev. Lett. 129, 238101 (2022).

[17] C. Mirabello, B. Wallner, B. Nystedt, S. Azinas, and M. Carroni, Unmasking AlphaFold to integrate experiments and predictions in multimeric complexes, Nat. Commun. 15, 8724 (2024).

[18] R. Yin and B. G. Pierce, Evaluation of AlphaFold antibody-antigen modeling with implications for improving predictive accuracy, Protein Sci. 33, e4865 (2024).

[19] Y. Lipman, R. T. Q. Chen, H. Ben-Hamu, M. Nickel, and M. Le, Flow matching for generative modeling, arXiv [cs.LG] (2022).

[20] G. Ahdritz, N. Bouatta, C. Floristean, S. Kadyan, Q. Xia, W. Gerecke, T. J. O’Donnell, D. Berenberg, I. Fisk, N. Zanichelli, B. Zhang, A. Nowaczynski, B. Wang, M. M. Stepniewska-Dziubinska, S. Zhang, A. Ojewole, M. E. Guney, S. Biderman, A. M. Watkins, S. Ra, P. R. Lorenzo, L. Nivon, B. Weitzner, Y.-E. A. Ban, S. Chen, M. Zhang, C. Li, S. L. Song, Y. He, P. K. Sorger, E. Mostaque, Z. Zhang, R. Bonneau, and M. AlQuraishi, OpenFold: retraining AlphaFold2 yields new insights into its learning mechanisms and capacity for generalization, Nat. Methods 21, 1514 (2024).

[21] D. Kovtun, M. Akdel, A. Goncearenco, G. Zhou, G. Holt, D. Baugher, D. Lin, Y. Adeshina, T. Castiglione, X. Wang, C. Marquet, M. McPartlon, T. Geffner, E. Rossi, G. Corso, H. Stärk, Z. Carpenter, E. Kucukbenli, M. Bronstein, and L. Naef, PINDER: The protein interaction dataset and evaluation resource, bioRxiv (2024).

[22] F. N. Hitawala and J. J. Gray, What does AlphaFold3 learn about antibody and nanobody docking, and what remains unsolved?, MAbs 17, 2545601 (2025).

[23] ESM Team, ESM cambrian: Revealing the mysteries of proteins with unsupervised learning, EvolutionaryScale Website (2024).

[24] W. Peebles and S. Xie, Scalable diffusion models with transformers, arXiv [cs.CV] (2022).

[25] M. S. Albergo and E. Vanden-Eijnden, Building normalizing flows with stochastic interpolants, arXiv [cs.LG] (2022).

[26] X. Liu, C. Gong, and Q. Liu, Flow straight and fast: Learning to generate and transfer data with rectified flow, arXiv [cs.LG] (2022).

[27] B. Jing, B. Berger, and T. Jaakkola, AlphaFold meets flow matching for generating protein ensembles, arXiv [q-bio.BM] (2024).

[28] Y. Yan, H. Tao, J. He, and S.-Y. Huang, The HDOCK server for integrated protein-protein docking, Nat. Protoc. 15, 1829 (2020).

[29] B. G. Pierce, K. Wiehe, H. Hwang, B.-H. Kim, T. Vreven, and Z. Weng, ZDOCK server: interactive docking prediction of protein-protein complexes and symmetric multimers, Bioinformatics 30, 1771 (2014).

[30] H. Hwang, T. Vreven, J. Janin, and Z. Weng, Protein-protein docking benchmark version 4.0, Proteins 78, 3111 (2010).

[31] R. F. Alford, A. Leaver-Fay, J. R. Jeliazkov, M. J. O’Meara, F. P. DiMaio, H. Park, M. V. Shapovalov, P. D. Renfrew, V. K. Mulligan, K. Kappel, J. W. Labonte, M. S. Pacella, R. Bonneau, P. Bradley, R. L. Dunbrack, Jr, R. Das, D. Baker, B. Kuhlman, T. Kortemme, and J. J. Gray, The rosetta all-atom energy function for macromolecular modeling and design, J. Chem. Theory Comput. 13, 3031 (2017).

[32] V. Stebliankin, A. Shirali, P. Baral, J. Shi, P. Chapagain, K. Mathee, and G. Narasimhan, Evaluating protein binding interfaces with transformer networks, Nat. Mach. Intell. 5, 1042 (2023).

[33] D. Kuroda and J. J. Gray, Pushing the backbone in protein-protein docking, Structure 24, 1821 (2016).

[34] A. Harmalkar and J. J. Gray, Advances to tackle backbone flexibility in protein docking, Curr. Opin. Struct. Biol. 67, 178 (2021).

[35] D. Del Alamo, D. Sala, H. S. Mchaourab, and J. Meiler, Sampling alternative conformational states of transporters and receptors with AlphaFold2, Elife 11 (2022).

[36] H. K. Wayment-Steele, A. Ojoawo, R. Otten, J. M. Apitz, W. Pitsawong, M. Hömberger, S. Ovchinnikov, L. Colwell, and D. Kern, Predicting multiple conformations via sequence clustering and AlphaFold2, Nature 625, 832 (2024).

[37] R. L. Dunbrack, Jr, Rēs ipSAE loquunt: What’s wrong with AlphaFold’s ipTM score and how to fix it, bioRxiv (2025).

[38] A. Harmalkar, S. Lyskov, and J. J. Gray, Reliable protein–protein docking with AlphaFold, rosetta, and replica exchange, Elife 13 (2025).

[39] D. Xu, L.-S. Chu, and J. Gray, Model weights and input structures for “adapting co-folding models for structure-based protein-protein docking through flow matching”, 10.5281/ZENODO.17782958 (2025).

