## Supplemental Material for "Adapting Co-Folding Models for Structure-Based Protein-Protein Docking Through Flow Matching"

April 23, 2026

#### 1 Additional details on training data

We subsample the PINDER training set similar to the original work [1]. We first perform a pre-filtering process where we remove entries that: (1) have more than one consecutive unknown residue (X) in their sequence; (2) have inconsistency in length between their sequence and residue numbers; (3) have a buried solvent-accessible surface area value lower than 400 Å$^2$; (4) are released after 2021-09-30 (so that we can directly compare our results with AF-M and AF3). We then additionally filter entries that: (1) have fewer than four atom types (low quality structures); (2) are too elongated (max_var>0.98); (3) have a sequence length shorter than 40 residues; (4) have a resolution worse than 5 Å. We cluster the remaining entries according to both their interface structure clusters (provided in PINDER) and their sequence clusters. The sequence clusters were obtained by running mmseqs2 easy-cluster command with a minimum sequence identity of 0.4 and a coverage of 0.5 on all protein sequences in the PINDER dataset. We pick the top two entries from each structure-sequence cluster, based on their apo and predicted structure availability, whether they were determined by X-ray crystallography, and their resolution. Due to other issues such as inconsistencies between the number of residues in the structure and residue numbers, as well as training samples being too large to fit into the RAM of our setup, we additionally remove 90 samples from the training set. This subsampling process finally results in 34,838 training samples of bi-protein complexes, with 9,088 samples containing at least one chain that has an apo structure, and 30,730 samples containing at least one chain that has a predicted structure.

In addition to bi-protein complexes in the PINDER dataset, we also construct tri-protein complexes as part of our training set. We find all groups of three protein chains that are interacting with each other in the PINDER training set, and then filter and subsample them based on the same criteria as the bi-protein case, except that we only pick the top entry from each structure-sequence cluster. During training, we randomly combine two chains in each three-chain group into a single chain with an added residue index gap of 200 between chains and use it as the receptor, while leaving the third chain as the ligand. This results in 10,515 tri-protein complexes, with 2,038 complexes containing at least one chain that has an apo structure, and 9,060 complexes containing at least one chain that has a predicted structure.

#### 2 Additional details on model training and inference of AF2Dock

**Model training.** We implement AF2Dock based on the OpenFold implementation [2] of AF-M in PyTorch. We use all original AF-M loss functions for training except for the masked MSA loss [3, 4]. We also adjust the resolution threshold for computing TM and pLDDT losses from 3 Å to 4 Å. We use a $\sigma_{\rm tr}$ of 30 Å for the translational prior in both training and inference. We center the translational Gaussian distribution on the ligand during training to bias the training samples closer to the ground truth pose, while we center the distribution on the receptor during inference. We initialize the model weights with AF-M version 2.3 model 1 weights for AF-M components, and randomized weights for the docking module. We train the model in three stages. For stage one, we freeze the weights of AF-M components and only train the docking module. We only train with holo monomer structures for this stage, using a learning rate of 0.001, a warmup phase of 1000 steps, and a batch size of 64, for around 5000 steps. For stage two, we finetune the docking module while still keeping the weights of AF-M components frozen. In this stage, for each protein chain, we randomly sample holo, apo, and predicted monomer structures with a probability of 20%, 40%, and 40%, when available. We train with a learning rate of 0.0005 and a batch size of 32 for around 7000 steps. For stage three, we perform full-parameter finetuning of AF-M along with the docking module. We use the same monomer sampling scheme as stage two, a learning rate of 0.0005, and a batch size of 32 for around 3000 steps. We use a weight of 0.03 for violation loss during the first two stages, and a weight of 0.5 during stage three. We use a crop size of 384 for all stages, same as AF-M. We observed occasional training instability during stage one, which is the reason we decided to use only holo structures in this stage. We also found that restarting training from an earlier checkpoint with a different seed helped stabilize the loss. We trained all models on 4x NVIDIA A100 GPUs with a variable number of gradient accumulation steps to match the batch size. For ablation studies, we use the same hyperparameters and training schemes, except that we either omit the input of ESM embeddings to the docking module, or stage three full-parameter fine-tuning.

**Model inference.** Inference of AF2Dock is performed on a single NVIDIA A100 GPU. We use the deepspeed evoformer attention kernel available in OpenFold [2] during inference, which speeds up computation and reduces memory cost. By default, we integrate over 10 time steps for each sample along the learned velocity field. We sample 20 poses for the PINDER-AF2 benchmark and 40 poses for the antibody nanobody set. We rank the resulting predictions by their ipTM score. For non-holo structure inputs in the PINDER-AF2 benchmark, we truncate the structures to the same residues as holo structures. In the case of antibody-antigen complexes, we merge the antibody heavy and light chains into a single chain and dock it as a single entity during inference, using the same method as three-chain samples during training (combine the two chains with an added residue index gap of 200 between them). We perform boostrapping to estimate the uncertainty of predictions. We represent the success rates as the mean and 95% confidence interval of 10,000 bootstrap samples obtained by randomly sampling the original predicted structures with replacement. For the CDRH3 inpainting experiment, we remove CDRH3 from the predicted antibody and nanobody structures, as well as any residue in both antibody/nanobody and antigen structures with a pLDDT score lower than 70.

### 3 Details on baseline model inference

**Single-sequence AF-M with custom template inputs** We perform predictions of single-sequence AF-M with custom template inputs using a modified OpenFold, producing one predicted structure per AF-M model (five total). By default, we use the original recycling parameters (20 maximum recycling iterations with early stop), although we also performed predictions without recycling as an additional baseline. As part of the ablation studies, we additionally perform predictions with two variants of non-recycling single-sequence AF-M with either full-complex templates of noisy structures sampled from the same prior distribution as AF2Dock, or on top of that, including the same interpolation process as AF2Dock. For these two variants, we only use AF-M model 1 and sample different noisy initial structures.

**Co-folding AF-M and AF3** For co-folding AF-M, we use the local ColabFold implementation [5] with MSAs generated from the mmseqs MSA server. For AF3, we use the official implementation. For both co-folding AF-M and AF3, we set the template cutoff date to 2021-09-30. We use the standard data pipeline and do not input the same templates as the structure-based docking methods.

**DFMDock** DFMDock inference was performed with default parameters using 40 diffusion steps.

**DiffDock-PP** DiffDock-PP inference was performed with default parameters used for Docking Benchmark 5.5 test in the original work using 40 diffusion steps.

**HDock** HDock inference was performed with HDock-Lite version 1.1 using default parameters.

**ZDock** ZDock inference was performed with ZDock version 3.0.2 using default parameters. The `mark_sur` program was first used to mark the surfaces of individual subunits, before inputting into the `zdock` program for pose inference.

For all sampling-based baseline model predictions, including co-folding AF-M, AF3, DFMDock, DiffDock-PP, and the two variants of single-sequence AF-M, we match the total number of samples (five times the number of seeds in the case AF-M and AF3) to be the same as AF2Dock. For single-sequence AF-M, we produce five predictions (one per model weight). For grid-search-based HDock and ZDock, we perform the default search process but only include the top 20 and 40 ranked samples as oracle for the PINDER-AF2 benchmark and the antibody/nanobody test set. We rank the resulting predictions with the default ranking scores of the respective models, and with ipTM for single-sequence AF-M and its variants. We perform boostrapping for all sampling-based methods in the same way as AF2Dock. We do not estimate uncertainties for single-sequence AF-M, HDock, and ZDock, as there is no stochasticity involved in their docking process.

### 4 Ranking predictions with external scoring metrics

One of the key limitations of AF2Dock is its inability to properly rank its predicted structures. To address this issue, we explored ranking AF2Dock predictions with external scoring metrics and compared the results with the default approach that ranks only using the internal AF2Dock ipTM. We tested three external scores: Rosetta interface score $I_{sc}$ [6] (combined with `docking_local_refine`), Piston score [7] (a surface-based scoring method), and AF-M ipTM obtained by inputting the predicted structure as a template to AF-M in single-sequence mode (which we dub AFMRank, as it is similar to the AF2Rank approach [8]). For AFMRank, we include inter-chain information in the input template, same as AlphaFold-Unmasked [9], and we only use the AF-M model 1 weight to perform the evaluation with no recycling. As the output of AF-M may differ from the input template structure, we additionally included an adjusted AFMRank score, which is the AF-M ipTM multiplied by the DockQ score between input and output structures, again following AF2Rank. We perform the ranking test on AF2Dock predicted structures for the antibody/nanobody test set using predicted inputs.

When we plot the scoring metrics against the DockQ scores of all predicted structures in the test set (Fig. S15), we observe that all scores tested here demonstrate the ability to discern good versus bad structures, as evidenced by the improved score mean going from low to high docking quality groups. However, all scoring metrics produce false positives on an absolute scale, assigning good scores to many samples with low DockQ. When pairs of scoring metrics are plotted against each other (Fig. S16), we observe a strong correlation between AF2Dock and AFMRank ipTM (Spearman $|\rho| \approx 0.7$), likely resulting from AF2Dock being fine-tuned from AF-M. Interestingly, the Rosetta $I_{sc}$ shows moderate correlation with AF2Dock and AFMRank ipTM scores (Spearman $|\rho|$ between 0.24 and 0.32). On the other hand, the Piston score exhibits the lowest correlation with other scores (Spearman $|\rho| \leq 0.15$).

When we compare the ranking success rates between different metrics (Fig. S17), we found that all metrics tested here produced comparable success rates. For antibody-antigen complexes, AF2Dock and AFMRank ipTM show the best performance, while for nanobody-antigen complexes, Rosetta $I_{sc}$ performs the best. However, when we plot the DockQ of Top-1 and Top-5 ranked structures by each scoring metrics against each other (Fig. S18 - S21), we found that each score successfully ranks a different set of targets, evidenced by the presence of targets in the top-left and bottom-right quadrants of the plots, similar to the orthogonality that we observed between different docking methods. The piston score shows particularly strong orthogonal ranking with the rest of metrics, consistent with its low correlations with others. The adjusted AFMRank ipTM did not result in better ranking compared to the unadjusted AFMRank ipTM in general. It is perhaps because the additional refinement performed by AF-M helps better identify whether the input is in the vicinity of a plausible configuration, and adjusting the score by downscaling highly confident yet different structures may hurt ranking performance. The orthogonality between scoring metrics suggests that using a combination of scores in ranking may lead to better results. Future work may rigorously explore this avenue by training and testing classification or ranking models that take various confidence metrics as inputs on a large set of predictions.

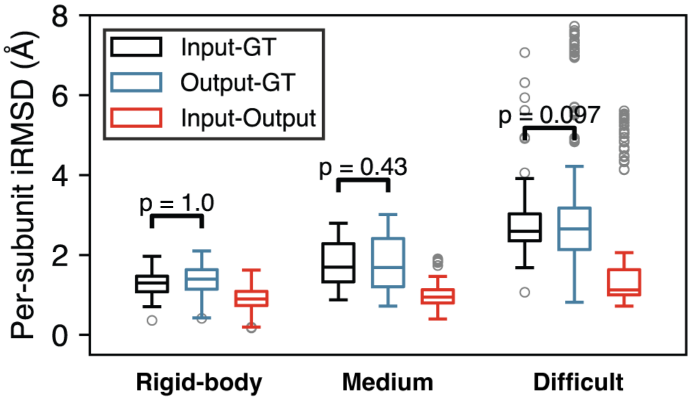


FIG. S1. Per-subunit interface RMSD (ps-iRMSD) among input, output, and ground truth (GT) structures for successful predictions in different difficulty categories of AF2Dock on the PINDER-AF2 predicted set. One-sided Wilcoxon signed-rank test was performed between input-GT and output-GT ps-iRMSD.

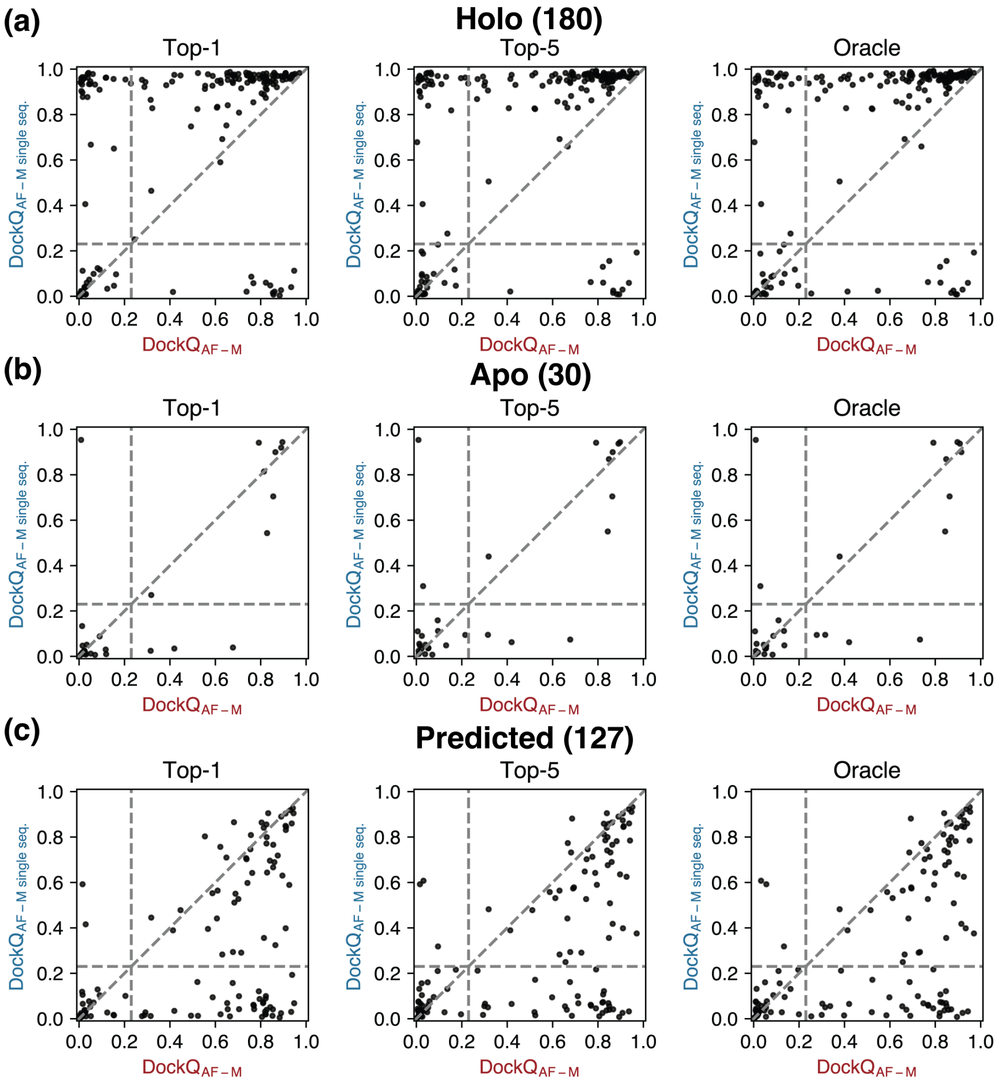


FIG. S2. Highest DockQ scores from the Top-1, Top-5, and oracle categories of predictions by single-sequence AF-M and co-folding AF-M plotted against each other for the PINDER-AF2 benchmark when using (a) holo, (b) apo, and (c) predicted structures in docking. Vertical and horizontal lines at DockQ $= 0.23$ (acceptable docking quality threshold), as well as the line $\mathrm{DockQ}_{\text{AF-M single seq.}} = \mathrm{DockQ}_{\text{AF-M}}$ are shown in gray as visual guides.

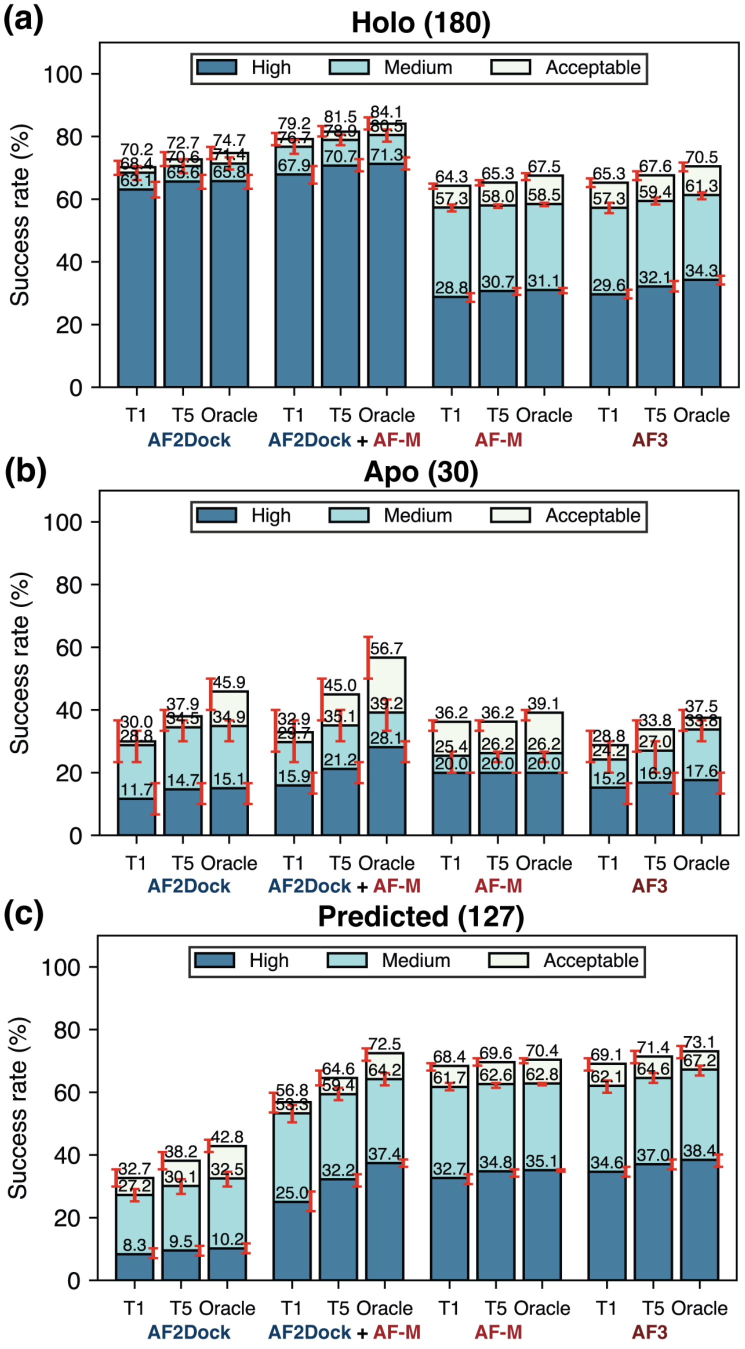


FIG. S3. Top-1 (T1), Top-5 (T5), and oracle success rates of AF2Dock, combination of AF2Dock and co-folding AF-M followed by re-ranking with ipTM, as well as co-folding AF-M and AF3 on the PINDER-AF2 benchmark when using (a) holo, (b) apo, and (c) predicted monomer structures in docking. The oracle category includes 20 predictions for all methods. For combination of AF2Dock and co-folding AF-M, we randomly sample 10 predictions from each method with replacement and combine. Bar heights and error bars represent the mean and the 95% confidence interval of 10,000 bootstrap samples. Docking quality is defined by DockQ thresholds, with acceptable > 0.23, medium > 0.49, and high > 0.80.

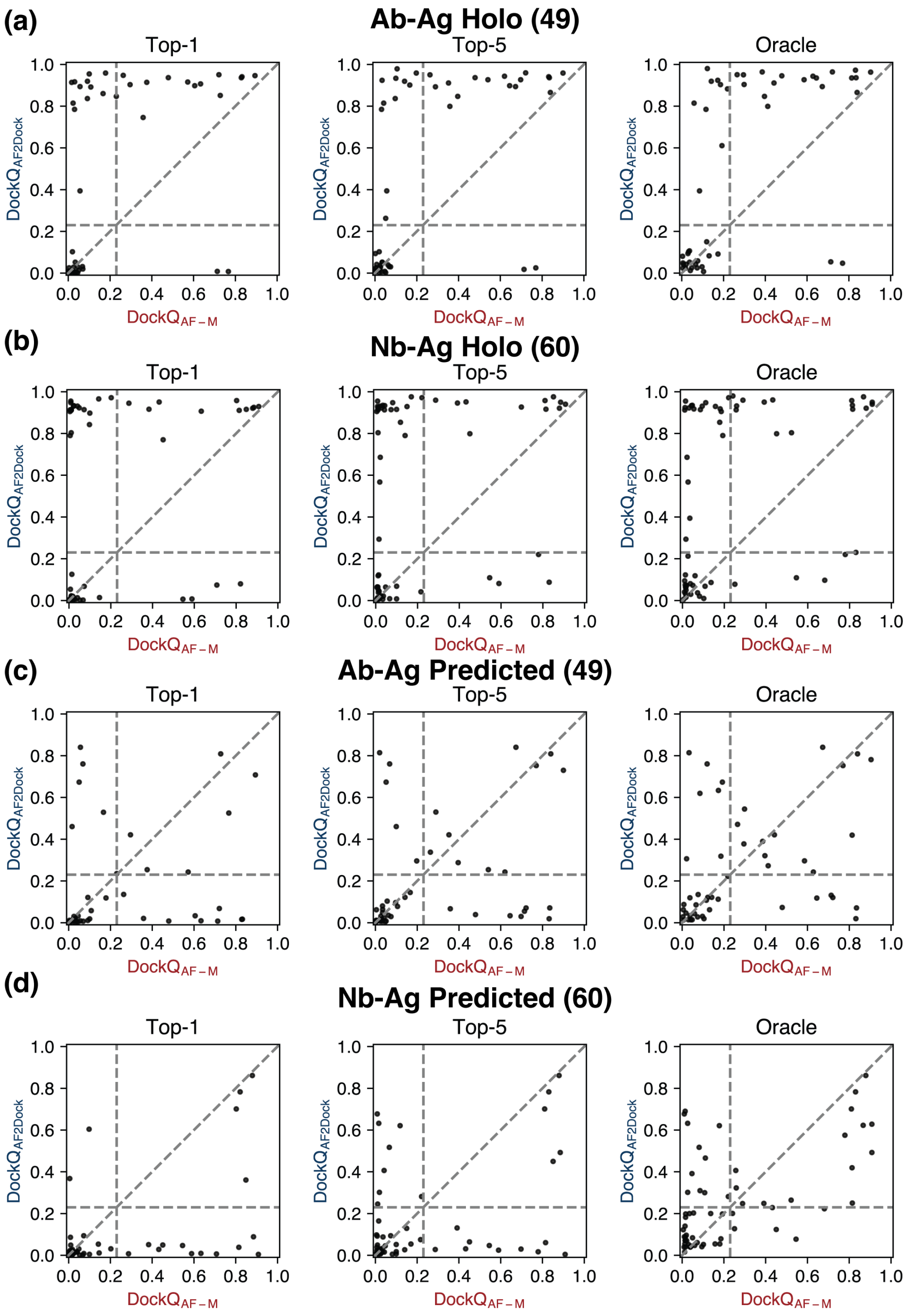


FIG. S4. Highest DockQ scores from the Top-1, Top-5, and oracle categories of predictions by AF2Dock and co-folding AF-M plotted against each other for antibody-antigen (Ab-Ag) complexes using (a) holo and (c) predicted inputs in docking, as well as for nanobody-antigen (Nb-Ag) complexes using (b) holo and (d) predicted inputs in docking. Vertical and horizontal lines at DockQ = 0.23 (acceptable docking quality threshold), as well as the line $\mathrm{DockQ_{AF2Dock}} = \mathrm{DockQ_{AF\text{-}M}}$ are shown in gray as visual guides.

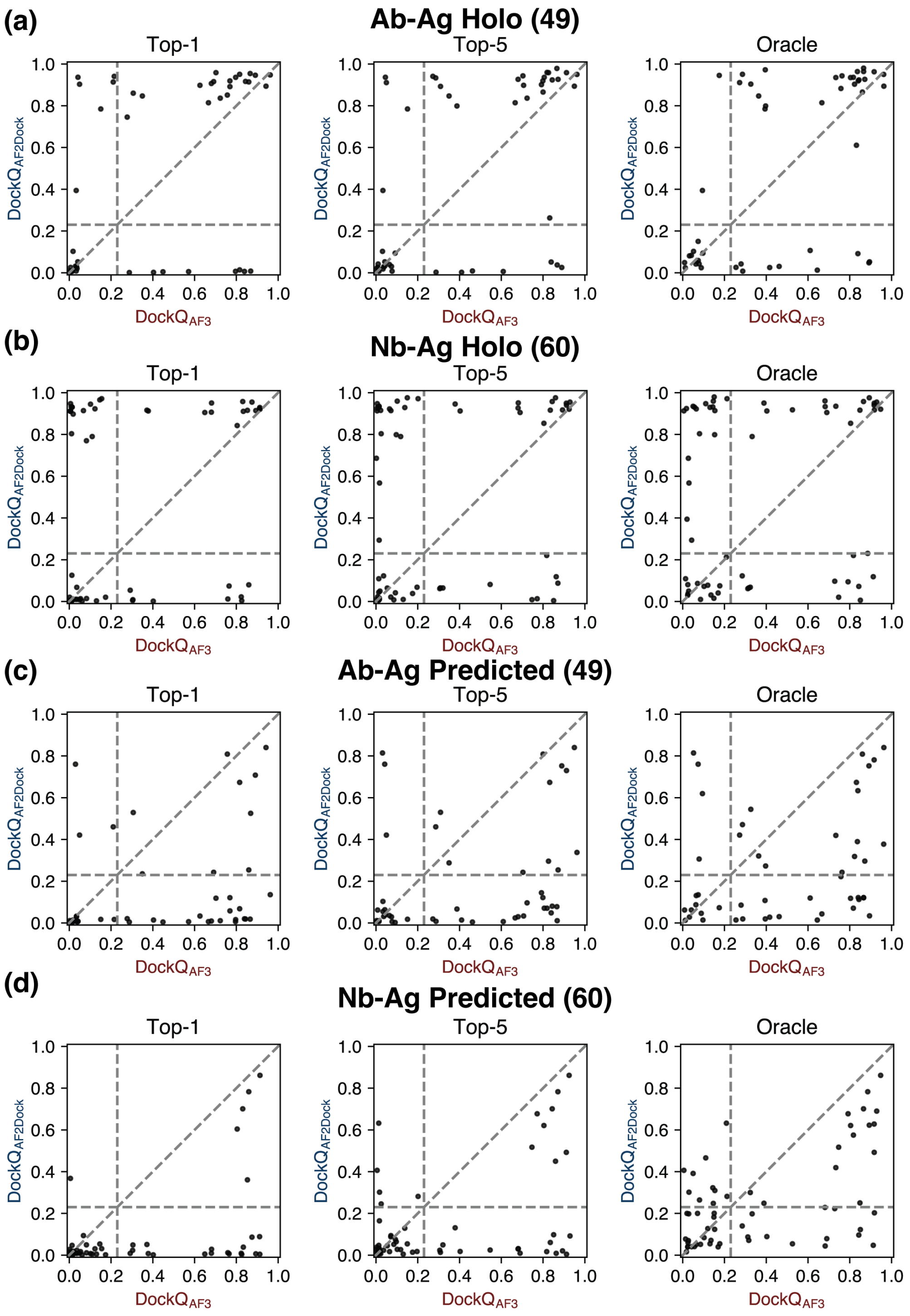


FIG. S5. Highest DockQ scores from the Top-1, Top-5, and oracle categories of predictions by AF2Dock and AF3 plotted against each other for antibody-antigen (Ab-Ag) complexes using (a) holo and (c) predicted inputs in docking, as well as for nanobody-antigen (Nb-Ag) complexes using (b) holo and (d) predicted inputs in docking. Vertical and horizontal lines at DockQ $= 0.23$ (acceptable docking quality threshold), as well as the line $\text{DockQ}_{\text{AF2Dock}} = \text{DockQ}_{\text{AF3}}$ are shown in gray as visual guides.

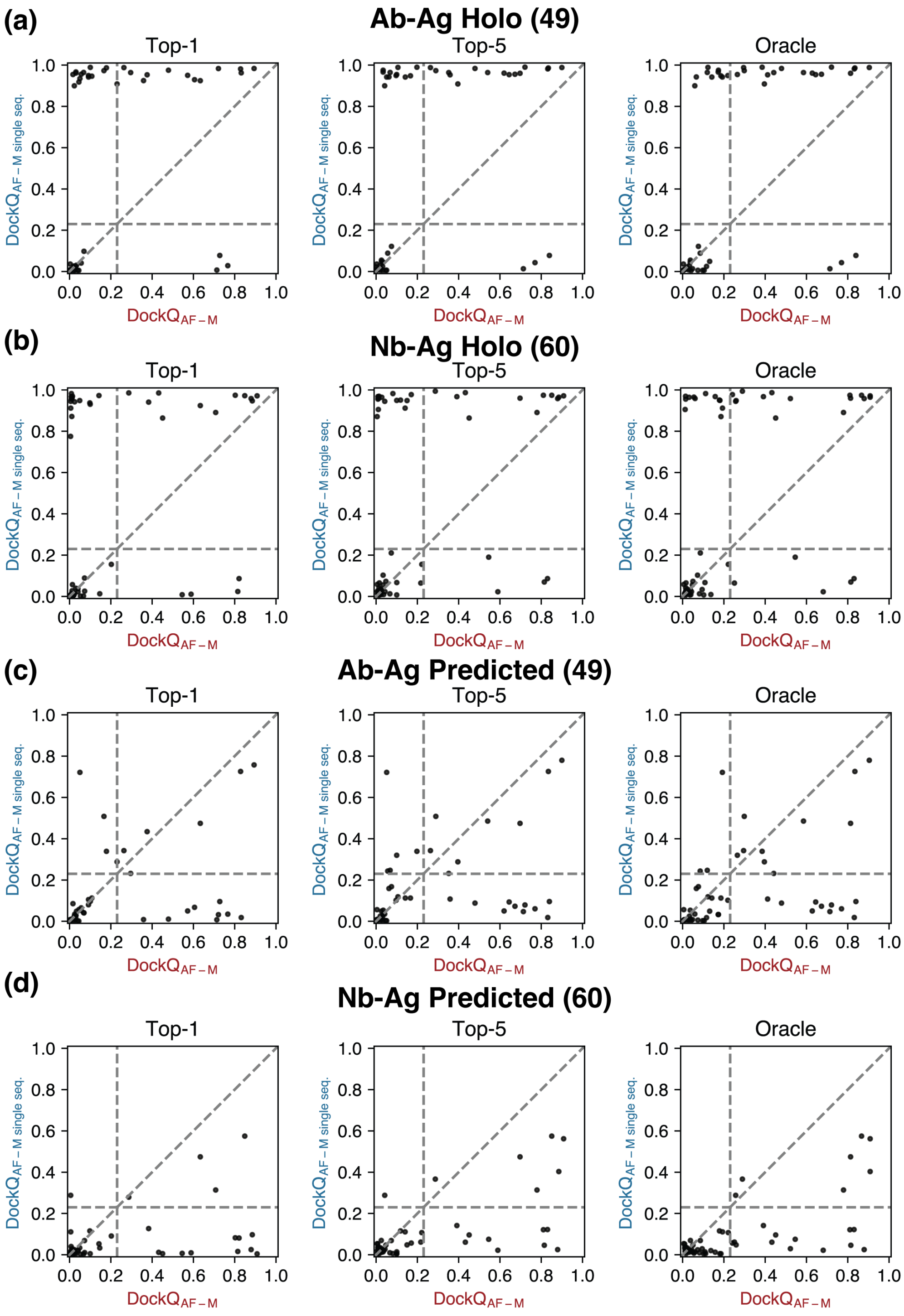


FIG. S6. Highest DockQ scores from the Top-1, Top-5, and oracle categories of predictions by single-sequence AF-M and co-folding AF-M plotted against each other for antibody-antigen (Ab-Ag) complexes using (a) holo and (c) predicted inputs in docking, as well as for nanobody-antigen (Nb-Ag) complexes using (b) holo and (d) predicted inputs in docking. Vertical and horizontal lines at DockQ = 0.23 (acceptable docking quality threshold), as well as the line $\text{DockQ}_{\text{AF-M single seq.}} = \text{DockQ}_{\text{AF-M}}$ are shown in gray as visual guides.

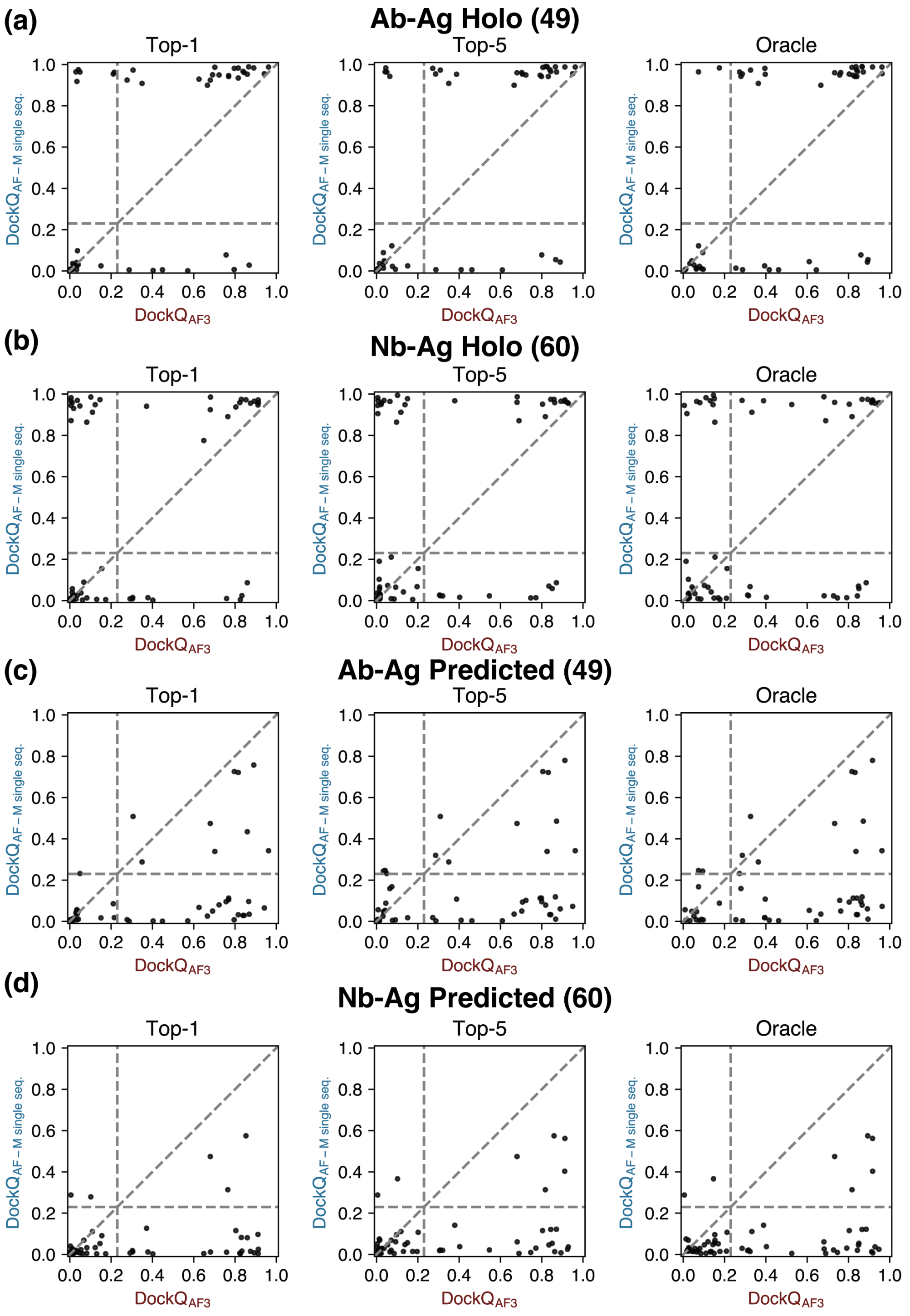


FIG. S7. Highest DockQ scores from the Top-1, Top-5, and oracle categories of predictions by single-sequence AF-M and AF3 plotted against each other for antibody-antigen (Ab-Ag) complexes using (a) holo and (c) predicted inputs in docking, as well as for nanobody-antigen (Nb-Ag) complexes using (b) holo and (d) predicted inputs in docking. Vertical and horizontal lines at DockQ = 0.23 (acceptable docking quality threshold), as well as the line DockQ$_{\text{AF-M single seq.}}$ = DockQ$_{\text{AF3}}$ are shown in gray as visual guides.

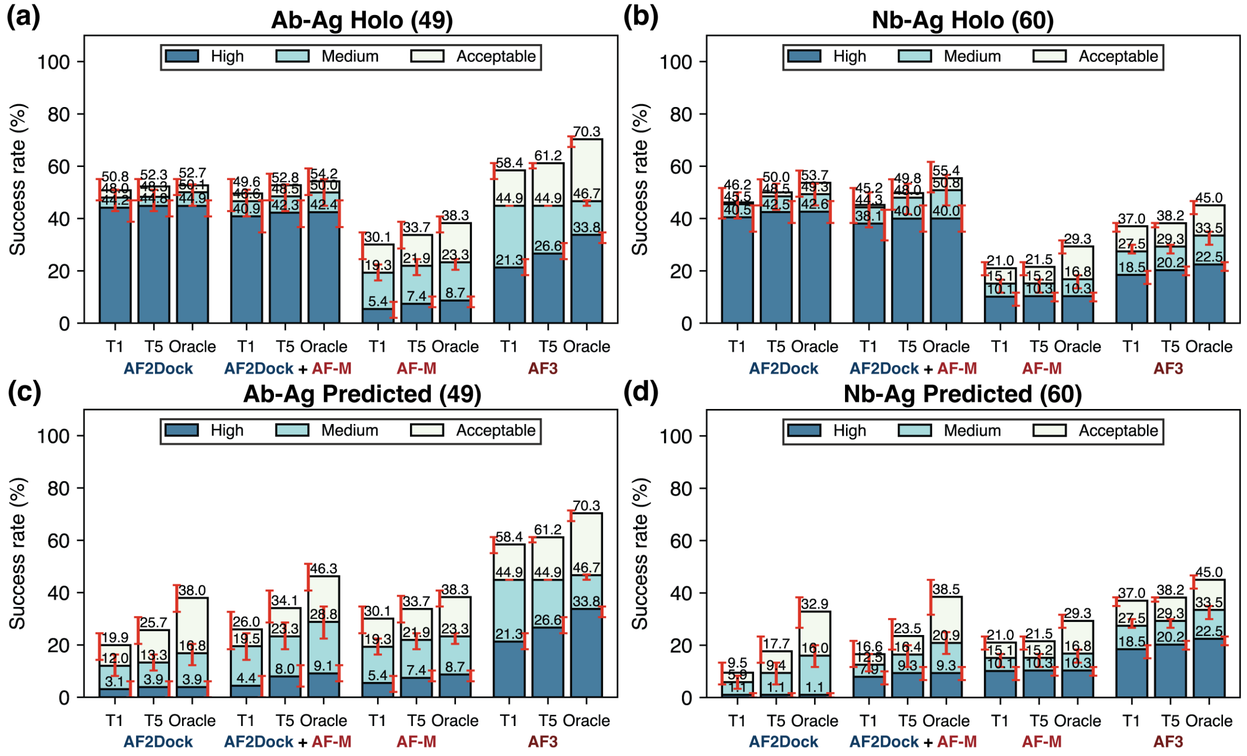


FIG. S8. Top-1 (T1), Top-5 (T5), and oracle success rates of AF2Dock, combination of AF2Dock and co-folding AF-M followed by re-ranking with ipTM, as well as co-folding AF-M and AF3 on (a,c) antibody-antigen (Ab-Ag) complexes and (b,d) nanobody-antigen (Nb-Ag) complexes when using (a,b) holo and (c,d) predicted inputs in docking. The oracle category includes 40 predictions for all methods. For combination of AF2Dock and co-folding AF-M, we randomly sample 20 predictions from each method with replacement and combine. Bar heights and error bars represent the mean and the 95% confidence interval of 10,000 bootstrap samples. Docking quality is defined by DockQ thresholds, with acceptable $> 0.23$, medium $> 0.49$, and high $> 0.80$.

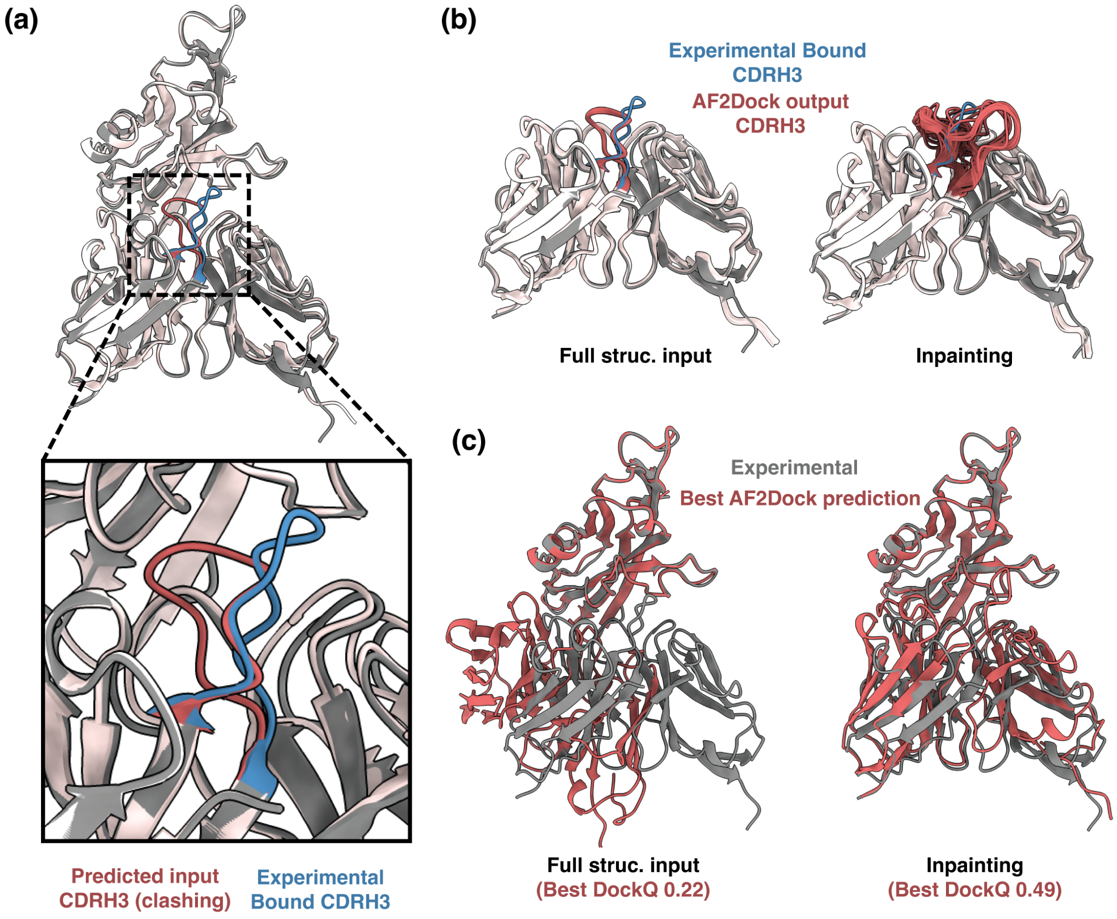


FIG. S9. An example case where inpainting CDRH3 and low pLDDT region lead to better predicitons (human VISTA extra cellular domain in complex with an antibody, PDB: 8tbq). (a) When predicted input structures are aligned with the experimental bound structure, the CDRH3 loop in the predicted structure (red) is in a very different conformation compared to the experimental structure (blue), which causes clashing. (b) Removing CDRH3 from input structures lead to much larger variability in the output CDRH3 conformations compared to using full predicted structure input. (c) Comparison of the output structure with the best DockQ score between using full predicted structure inputs and the inpainting strategy. The experimental structure is shown in gray and output structures are shown in red.

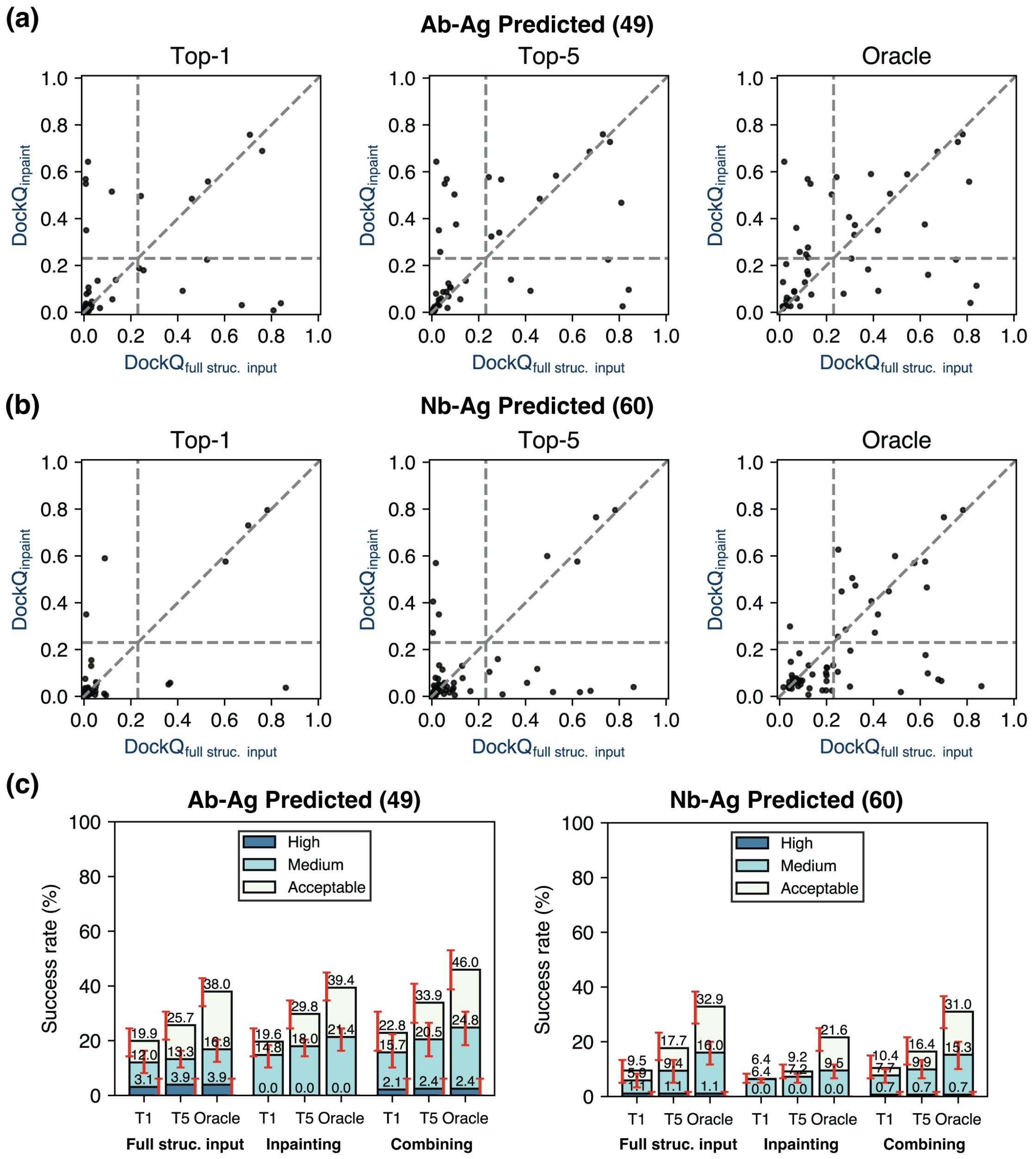


FIG. S10. Performance comparison between using full structural inputs and inpainting CDRH3. (a-b) Highest DockQ scores from the Top-1, Top-5, and oracle categories of predictions using full structural inputs and the inpainting strategy plotted against each other for (a) antibody-antigen (Ab-Ag) complexes and (b) nanobody-antigen (Nb-Ag) complexes when using predicted inputs in docking. Vertical and horizontal lines at DockQ = 0.23 (acceptable docking quality threshold), as well as the line $\text{DockQ}_{\text{inpaint}} = \text{DockQ}_{\text{full struc. input}}$ are shown in gray as visual guides. (c) Top-1 (T1), Top-5 (T5), and oracle success rates of AF2Dock on antibody-antigen (Ab-Ag) complexes (left) and nanobody-antigen (Nb-Ag) complexes (right) when using full predicted structure inputs, using input structures in which CDRH3 and low pLDDT regions are removed (inpainting), and when the predictions from the previous two approaches are combined and re-ranked with ipTM. The oracle category includes 40 predictions for all methods. For combination, we randomly sample 20 predictions from each approach with replacement and combine. Bar heights and error bars represent the mean and the 95% confidence interval of 10,000 bootstrap samples. Docking quality is defined by DockQ thresholds, with acceptable > 0.23, medium > 0.49, and high > 0.80.

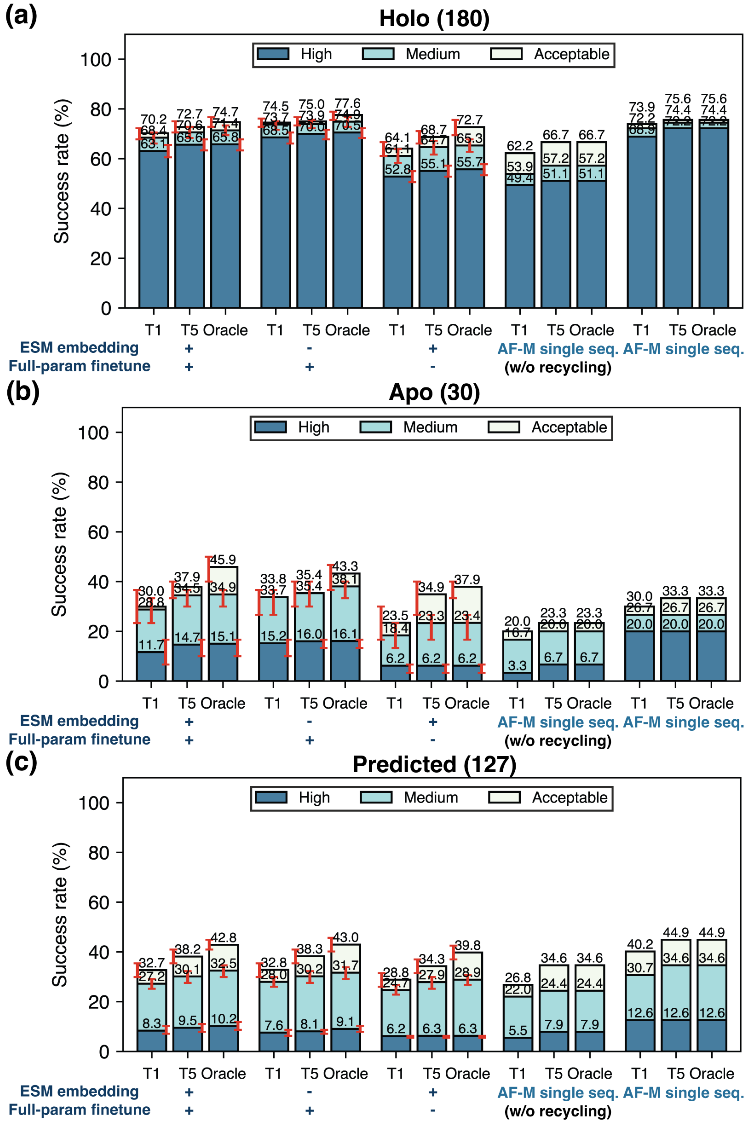


FIG. S11. Top-1 (T1), Top-5 (T5), and oracle success rates of base AF2Dock, two ablation variants of AF2Dock, and single-sequence AF-M on the PINDER-AF2 benchmark when using (a) holo, (b) apo, and (c) predicted monomer structures in docking. The oracle category includes 5 predictions for single-sequence AF-M and 20 predictions for other methods. Bar heights and error bars represent the mean and the 95% confidence interval of 10,000 bootstrap samples. We do not compute uncertainties for single-sequence AF-M, as there is no stochasticity in its sampling process. Docking quality is defined by DockQ thresholds, with acceptable $> 0.23$, medium $> 0.49$, and high $> 0.80$.

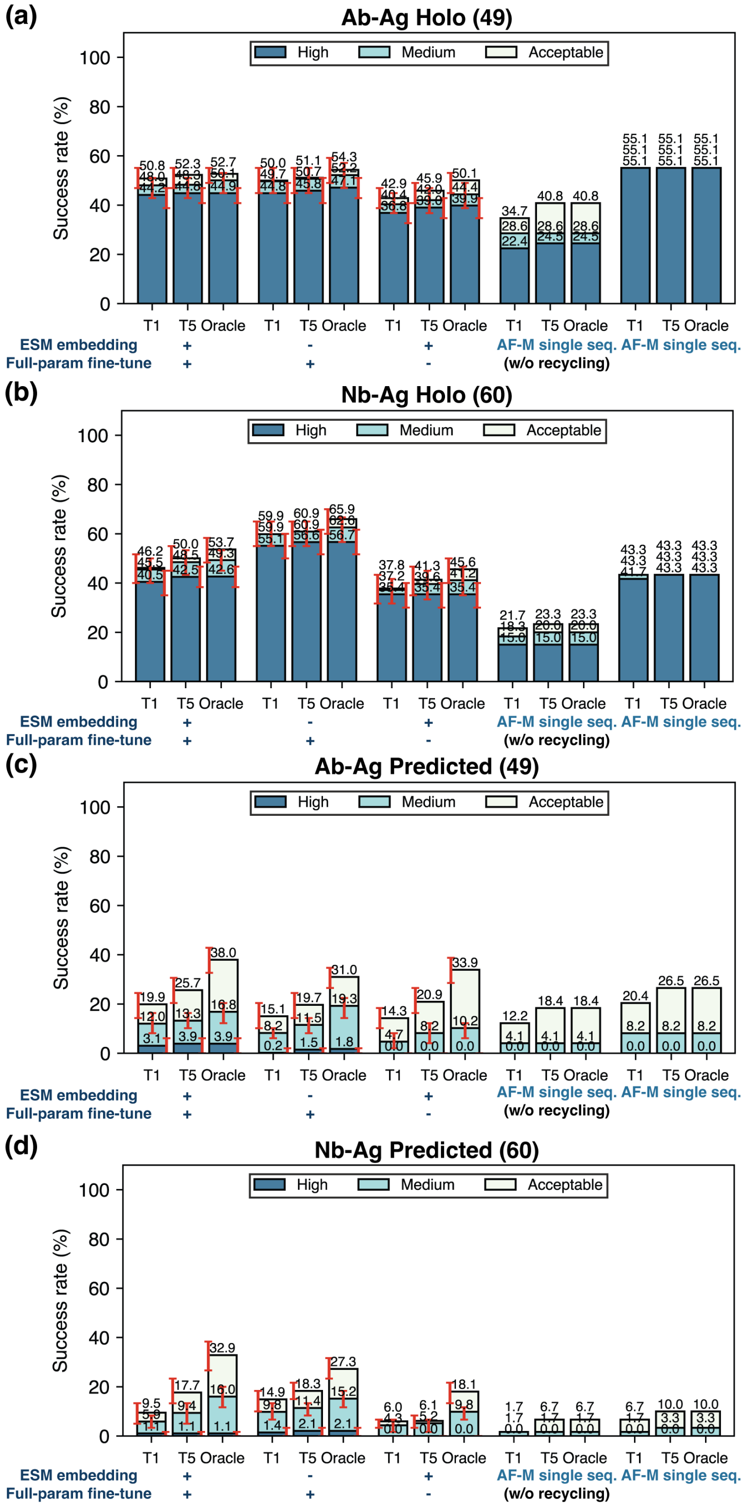


FIG. S12. Top-1 (T1), Top-5 (T5), and oracle success rates of base AF2Dock, two ablation variants of AF2Dock, and single-sequence AF-M on (a,c) antibody-antigen (Ab-Ag) complexes and (b,d) nanobody-antigen (Nb-Ag) complexes when using (a,b) holo and (c,d) predicted inputs in docking. The oracle category includes 5 predictions for single-sequence AF-M and 40 predictions for other methods. Bar heights and error bars represent the mean and the 95% confidence interval of 10,000 bootstrap samples. We do not compute uncertainties for single-sequence AF-M, as there is no stochasticity in its sampling process. Docking quality is defined by DockQ thresholds, with acceptable $> 0.23$, medium $> 0.49$, and high $> 0.80$.

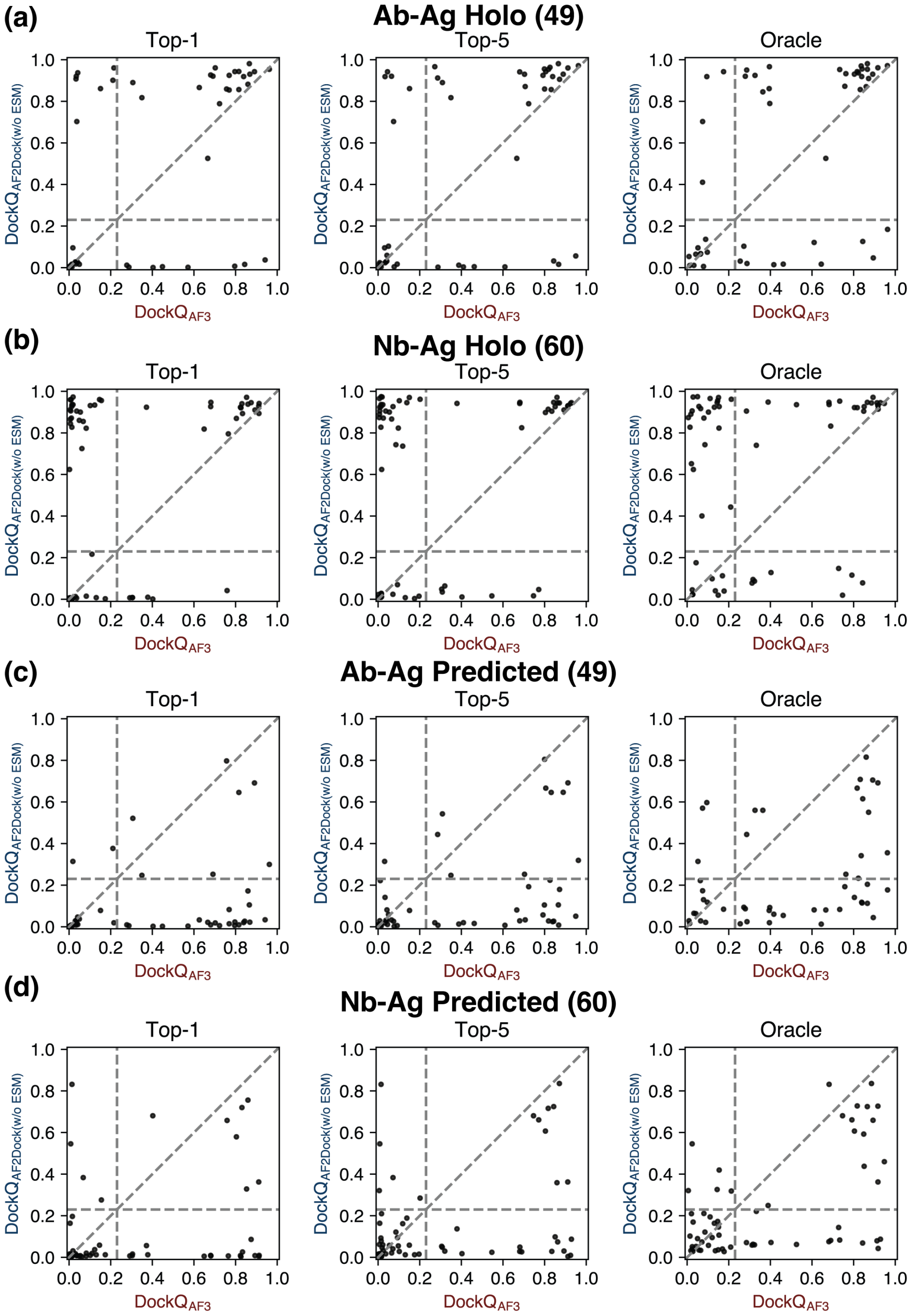


FIG. S13. Highest DockQ scores from the Top-1, Top-5, and oracle categories of predictions by the non-ESM variant of AF2Dock and AF3 plotted against each other for antibody-antigen (Ab-Ag) complexes when using (a) holo and (c) predicted inputs in docking, as well as for nanobody-antigen (Nb-Ag) complexes when using (b) holo and (d) predicted inputs in docking. Vertical and horizontal lines at DockQ = 0.23 (acceptable docking quality threshold), as well as the line $\mathrm{DockQ}_{\mathrm{AF2Dock(w/o\ ESM)}} = \mathrm{DockQ}_{\mathrm{AF3}}$ are shown in gray as visual guides.

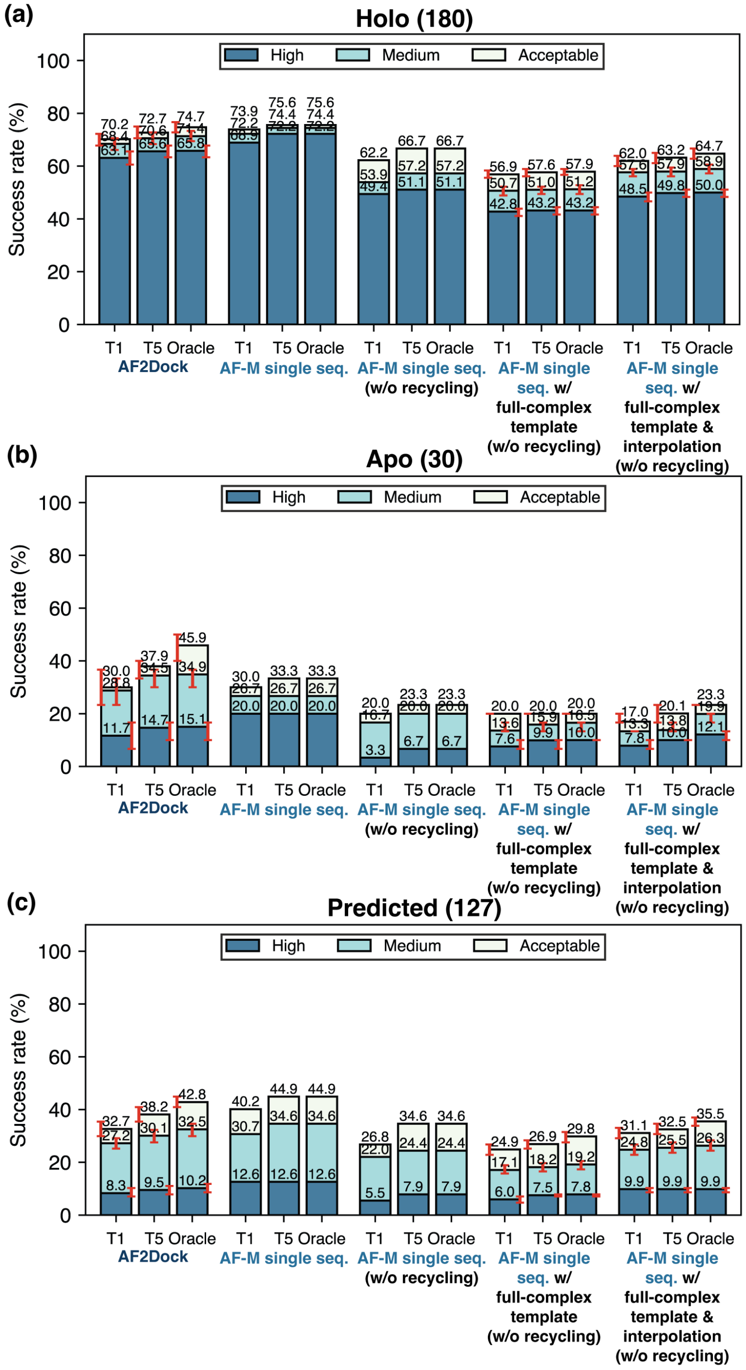


FIG. S14. Top-1 (T1), Top-5 (T5), and oracle success rates of AF2Dock, single-sequence AF-M, as well as two variants of single-sequence AF-M on the PINDER-AF2 benchmark when using (a) holo, (b) apo, and (c) predicted monomer structures in docking. The oracle category includes 5 predictions for single-sequence AF-M and 20 predictions for other methods. Bar heights and error bars represent the mean and the 95% confidence interval of 10,000 bootstrap samples. We do not compute uncertainties for vanilla single-sequence AF-M, as there is no stochasticity in its sampling process. Docking quality is defined by DockQ thresholds, with acceptable $> 0.23$, medium $> 0.49$, and high $> 0.80$.

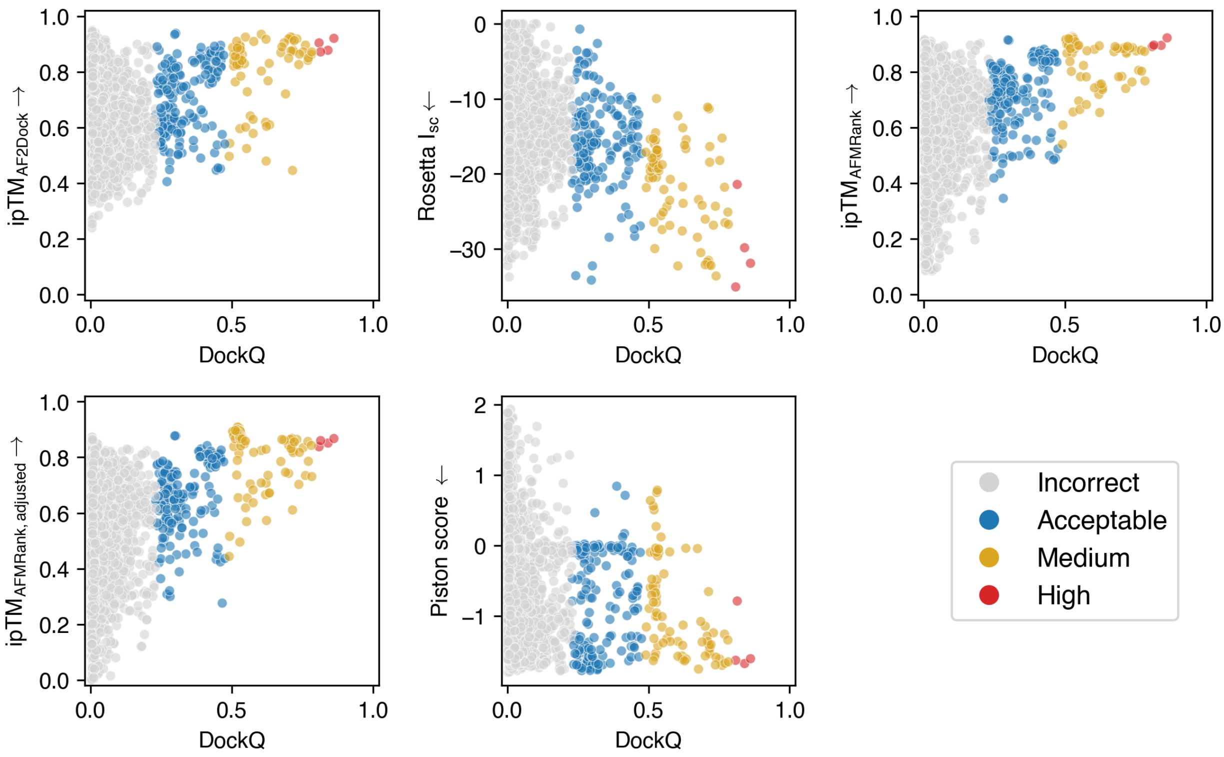


FIG. S15. Scoring metrics including AF2Dock ipTM, Rosetta interface score $I_{sc}$, AFMRank ipTM, adjusted AFMRank ipTM, and Piston score, plotted against DockQ on AF2Dock predicted structures for all targets in the antibody/nanobody test set using predicted input structures. Arrows on axis labels denote the direction towards a better score. Individual structures are colored by their docking quality, which is defined by DockQ thresholds, with incorrect $< 0.23$ (light gray), acceptable $> 0.23$ (blue), medium $> 0.49$ (gold), and high $>$ 0.80 (red).

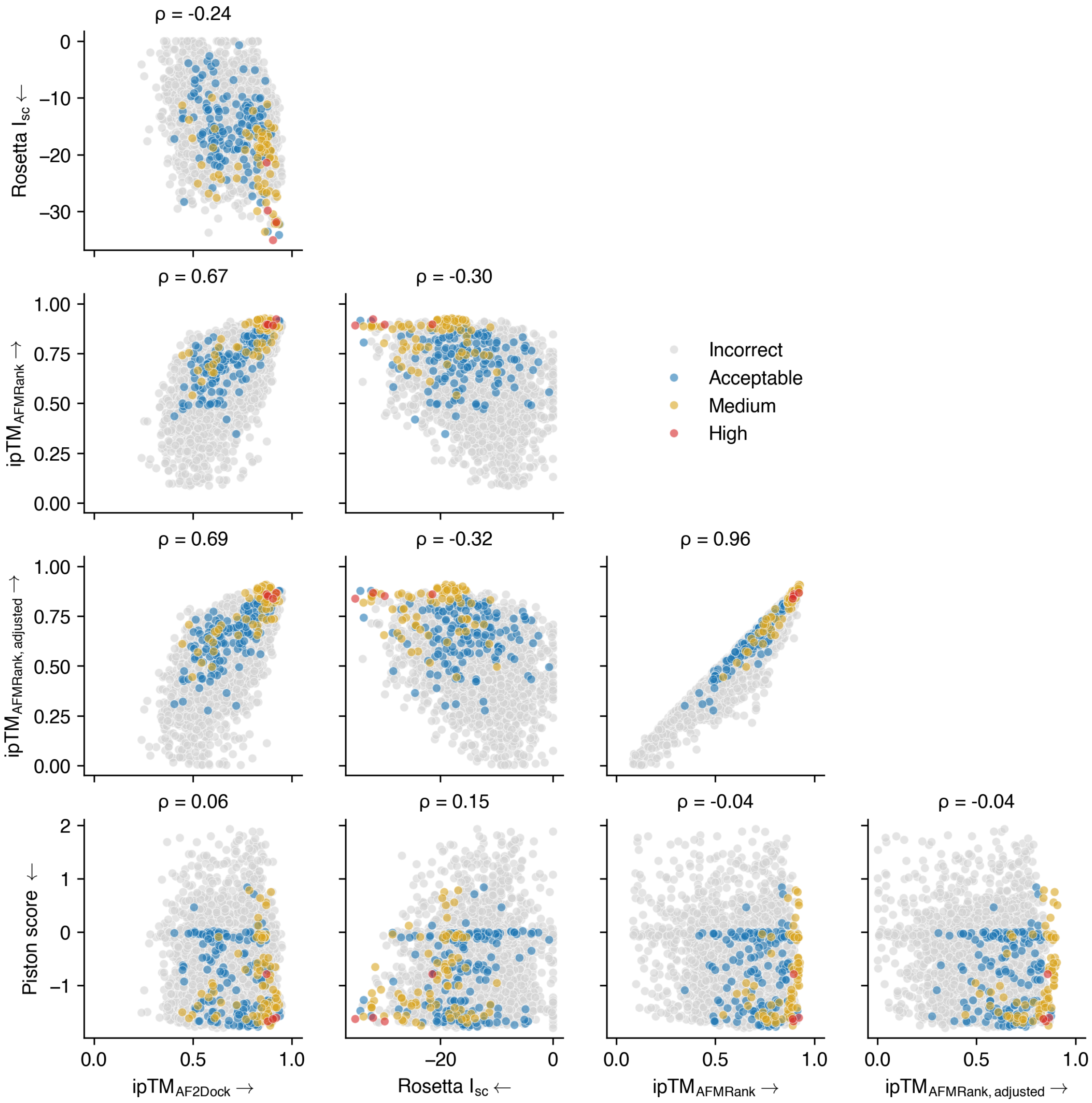


FIG. S16. Scoring metrics including AF2Dock ipTM, Rosetta interface score $I_{sc}$, AFMRank ipTM, adjusted AFMRank ipTM, and Piston score, plotted against each other on AF2Dock predicted structures for all targets in the antibody/nanobody test set using predicted input structures. Arrows on axis labels denote the direction towards a better score. Spearman correlations $\rho$ between metrics are computed and shown on top of each panel. Individual structures are colored by their docking quality, which is defined by DockQ thresholds, with incorrect $< 0.23$ (light gray), acceptable $> 0.23$ (blue), medium $> 0.49$ (gold), and high $> 0.80$ (red).

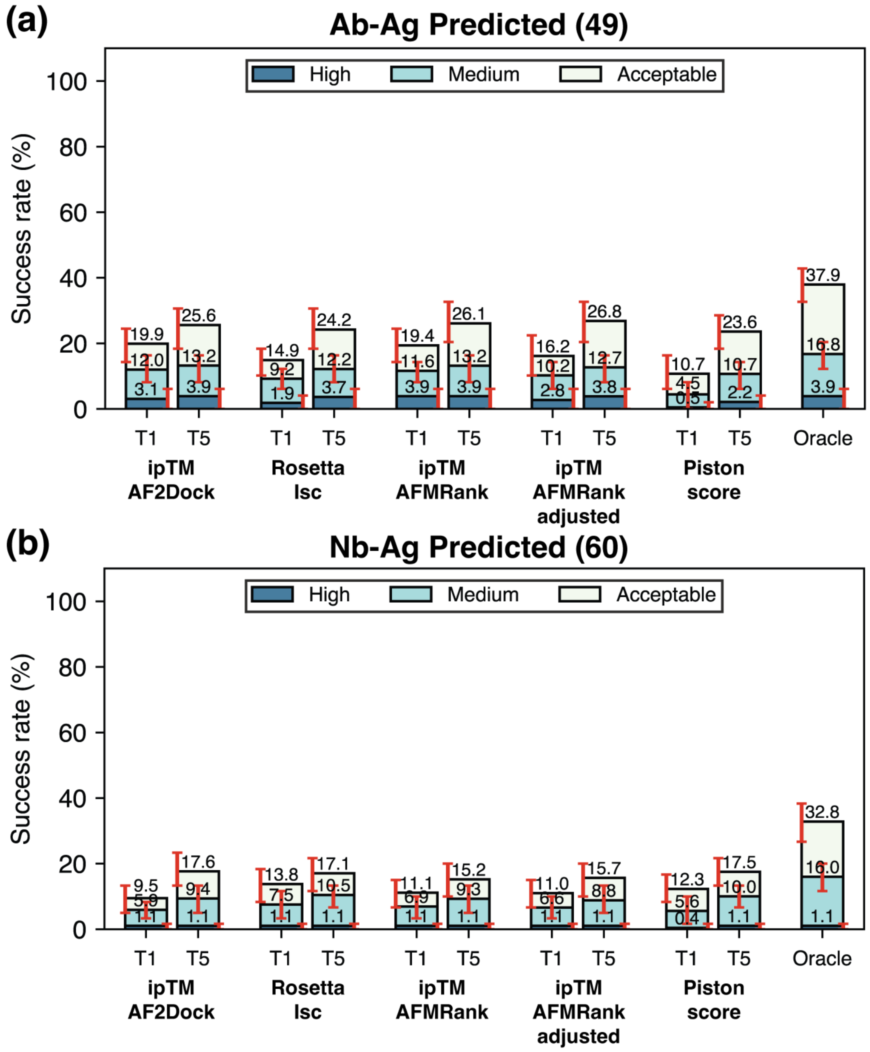


FIG. S17. Top-1 (T1) and Top-5 (T5) success rates from ranking with various scoring metrics on AF2Dock predictions for the antibody/nanobody test set using predicted input structures. Ranking was performed with AF2Dock ipTM, Rosetta interface score $I_{sc}$, AFMRank ipTM, adjusted AFMRank ipTM, and Piston score, on (a) antibody-antigen (Ab-Ag) complexes as well as (b) nanobody-antigen (Nb-Ag) complexes. The oracle success rate is identical for all cases and is shown on the far right. Bar heights and error bars represent the mean and the 95% confidence interval of 10,000 bootstrap samples. Docking quality is defined by DockQ thresholds, with acceptable > 0.23, medium > 0.49, and high > 0.80.

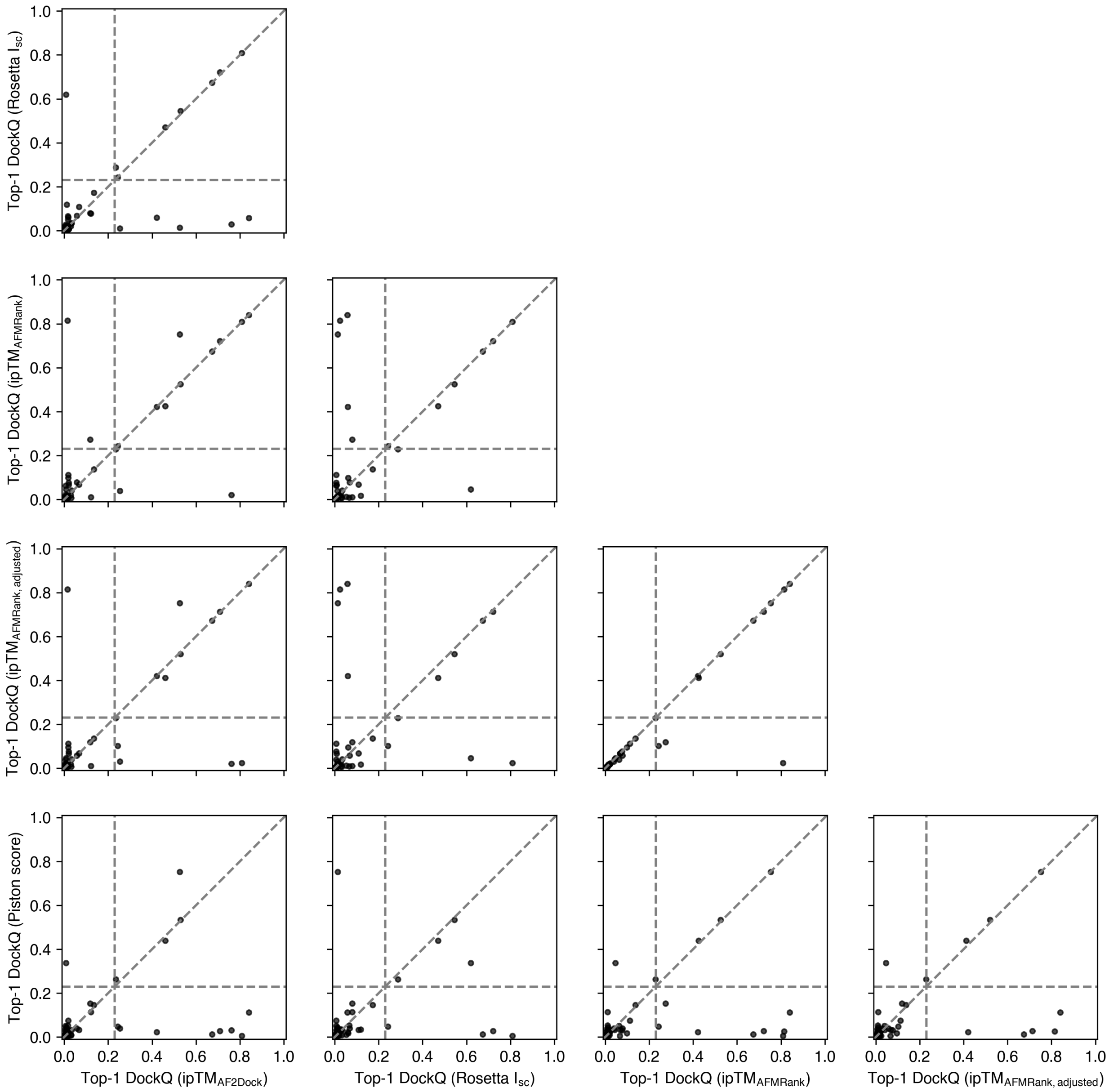


FIG. S18. Top-1 DockQ scores of AF2Dock predictions ranked by various scoring metrics plotted against each other for antibody-antigen (Ab-Ag) complexes using predicted input structures. Ranking was performed with AF2Dock ipTM, Rosetta interface score $I_{sc}$, AFMRank ipTM, adjusted AFMRank ipTM, and Piston score. Vertical and horizontal lines at DockQ = 0.23 (acceptable docking quality threshold), as well as the line $\mathrm{DockQ}_{\mathrm{score1}} = \mathrm{DockQ}_{\mathrm{score2}}$ are shown in gray as visual guides.

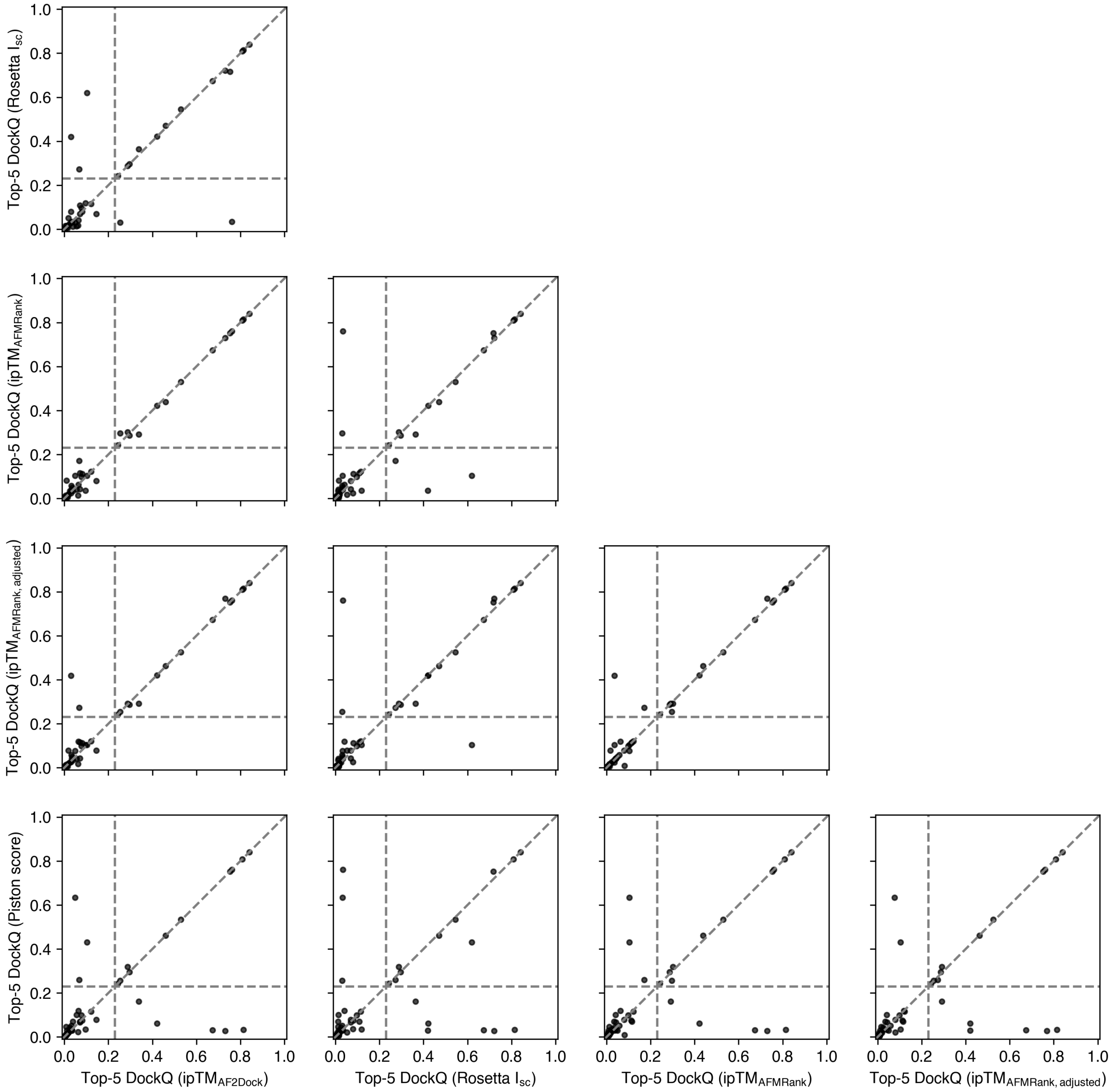


FIG. S19. Highest DockQ scores from the Top-5 category of AF2Dock predictions ranked by various scoring metrics plotted against each other for antibody-antigen (Ab-Ag) complexes using predicted input structures. Ranking was performed with AF2Dock ipTM, Rosetta interface score $I_{sc}$, AFMRank ipTM, adjusted AFM-Rank ipTM, and Piston score. Vertical and horizontal lines at DockQ = 0.23 (acceptable docking quality threshold), as well as the line $\mathrm{DockQ}_{\mathrm{score1}} = \mathrm{DockQ}_{\mathrm{score2}}$ are shown in gray as visual guides.

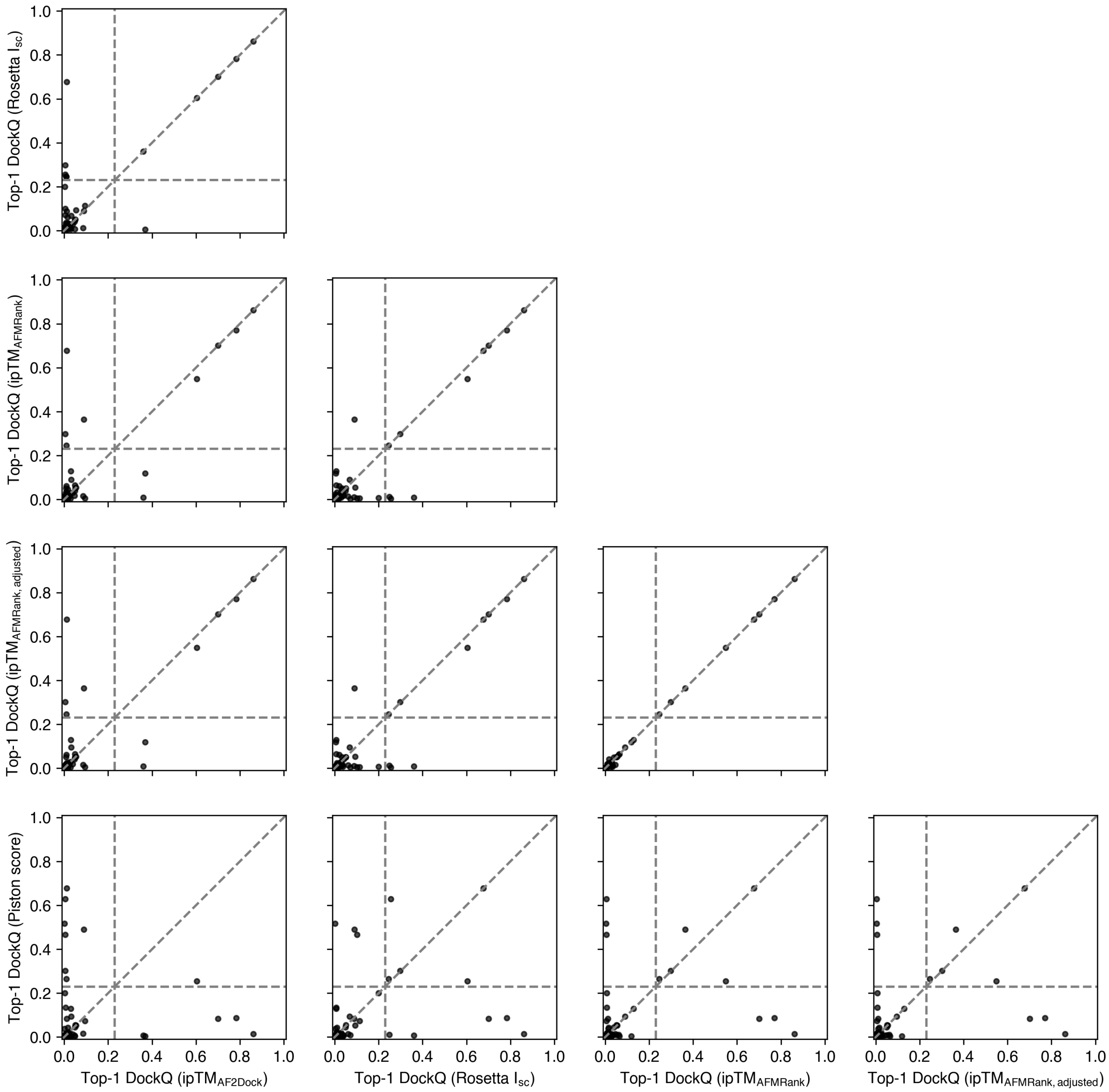


FIG. S20. Top-1 DockQ scores of AF2Dock predictions ranked by various scoring metrics plotted against each other for nanobody-antigen (Nb-Ag) complexes using predicted input structures. Ranking was performed with AF2Dock ipTM, Rosetta interface score $I_{sc}$, AFMRank ipTM, adjusted AFMRank ipTM, and Piston score. Vertical and horizontal lines at DockQ = 0.23 (acceptable docking quality threshold), as well as the line $\text{DockQ}_{\text{score1}} = \text{DockQ}_{\text{score2}}$ are shown in gray as visual guides.

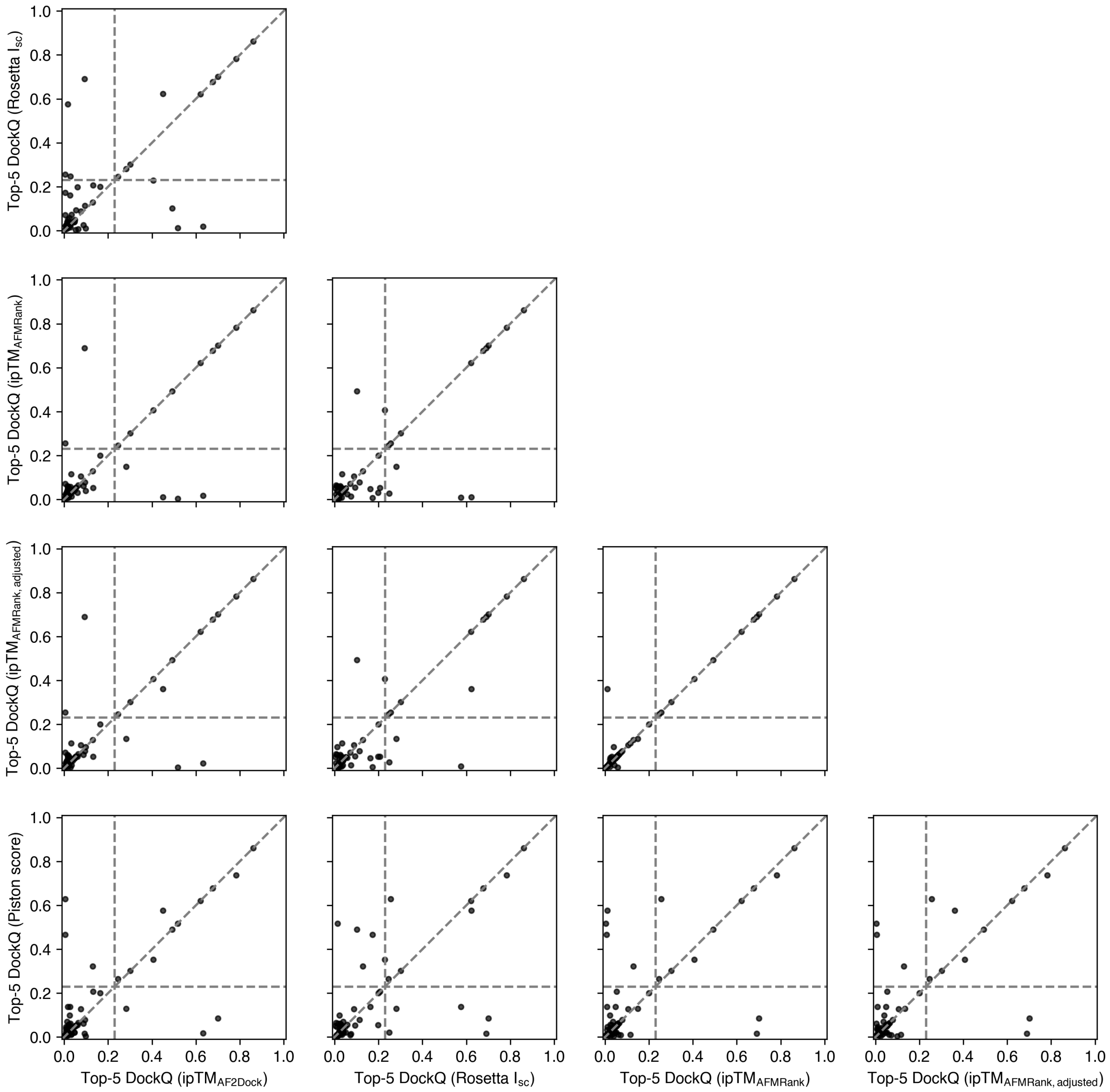


FIG. S21. Highest DockQ scores from the Top-5 category of AF2Dock predictions ranked by various scoring metrics plotted against each other for nanobody-antigen (Nb-Ag) complexes using predicted input structures. Ranking was performed with AF2Dock ipTM, Rosetta interface score $I_{sc}$, AFMRank ipTM, adjusted AFMRank ipTM, and Piston score. Vertical and horizontal lines at DockQ = 0.23 (acceptable docking quality threshold), as well as the line $\mathrm{DockQ}_{\mathrm{score1}} = \mathrm{DockQ}_{\mathrm{score2}}$ are shown in gray as visual guides.

**Algorithm S1** Structure embedder

**Input:** Noisy structure $\mathbf{x}_t$, sequence $a$, ESM embedding $\mathcal{E}$, pair representation $\mathbf{z}$
**Output:** Noisy pair representation update $\mathbf{u}$

**function** structure_embedder($\mathbf{x}_t, \{a_i\}, \{\mathcal{E}_i\}, \{\mathbf{z}_{ij}\},$ pair repr. latent dimension $c = 128$)
  $\{\mathbf{f}_{ij}^{\text{distogram}}, \mathrm{f}_{ij}^{\text{backbone\_frame\_mask}}, \mathbf{f}_{ij}^{\text{unit\_vector}}, \mathrm{f}_{ij}^{\text{pseudo\_beta\_mask}}\} \leftarrow \text{Compute}(\mathbf{x}_t)$
  $\mathbf{f}_i^{\text{restype}} \leftarrow \text{OneHot}(a_i)$
  $\mathbf{b}_{ij} \leftarrow \text{Concat}(\mathbf{f}_{ij}^{\text{distogram}}, \mathrm{f}_{ij}^{\text{backbone\_frame\_mask}}, \mathbf{f}_{ij}^{\text{unit\_vector}}, \mathrm{f}_{ij}^{\text{pseudo\_beta\_mask}})$
  $\mathbf{b}_{ij} \leftarrow \text{Concat}(\mathbf{b}_{ij}, \mathbf{f}_i^{\text{restype}}, \mathbf{f}_j^{\text{restype}}, \mathcal{E}_i, \mathcal{E}_j)$
  $\mathbf{b}_{ij} \leftarrow \text{Linear}(\text{LayerNorm}(\mathbf{z}_{ij})) + \text{Linear}(\mathbf{b}_{ij})$ ▷ $\mathbf{b}_{ij} \in \mathbb{R}^c$
  $\{\mathbf{b}_{ij}\} \leftarrow \text{PairStack}(\{\mathbf{b}_{ij}\}, N_{block} = 2, c = 128, c_{\text{mul}} = 128, c_{\text{att}} = 32, N_{\text{head}} = 4)$
  $\mathbf{u}_{ij} \leftarrow \text{Linear}(\text{ReLU}(\mathbf{b}_{ij}))$ ▷ $\mathbf{u}_{ij} \in \mathbb{R}^c$
  **return** $\{\mathbf{u}_{ij}\}$
**end function**

**Algorithm S2** Conditioning embedder

**Input:** Noisy structure $\mathbf{x}_t$, sequence $a$, ESM embedding $\mathcal{E}$, pair representation $\mathbf{z}$, time $t$
**Output:** Pair conditioning $\mathbf{g}$

**function** conditioning_embedder($\mathbf{x}_t, \{a_i\}, \{\mathcal{E}_i\}, \{\mathbf{z}_{ij}\}, t,$ pair repr. latent dimension $c = 128$)
  $\{\mathbf{f}_{ij}^{\text{distogram}}, \mathrm{f}_{ij}^{\text{backbone\_frame\_mask}}, \mathbf{f}_{ij}^{\text{unit\_vector}}, \mathrm{f}_{ij}^{\text{pseudo\_beta\_mask}}\} \leftarrow \text{Compute}(\mathbf{x}_t)$
  $\mathbf{f}_i^{\text{restype}} \leftarrow \text{OneHot}(a_i)$
  $\mathbf{b}_{ij} \leftarrow \text{Concat}(\mathbf{f}_{ij}^{\text{distogram}}, \mathrm{f}_{ij}^{\text{backbone\_frame\_mask}}, \mathbf{f}_{ij}^{\text{unit\_vector}}, \mathrm{f}_{ij}^{\text{pseudo\_beta\_mask}})$
  $\mathbf{b}_{ij} \leftarrow \text{Concat}(\mathbf{b}_{ij}, \mathbf{f}_i^{\text{restype}}, \mathbf{f}_j^{\text{restype}}, \mathcal{E}_i, \mathcal{E}_j)$
  $\mathbf{b}_{ij} \leftarrow \text{Linear}(\text{LayerNorm}(\mathbf{z}_{ij})) + \text{Linear}(\mathbf{b}_{ij})$ ▷ $\mathbf{b}_{ij} \in \mathbb{R}^c$
  $\mathbf{b}_{ij} \leftarrow \text{LayerNorm}(\mathbf{b}_{ij})$
  $\mathbf{b}_{ij} \leftarrow \mathbf{b}_{ij} + \text{LinearNoBias}(\text{LayerNorm}(\text{FourierEmbedding}(t, c = 256)))$
  $\{\mathbf{b}_{ij}\} \leftarrow \text{PairStack}(\{\mathbf{b}_{ij}\}, N_{\text{block}} = 2, c = 128, c_{\text{mul}} = 128, c_{\text{att}} = 32, N_{\text{head}} = 4)$
  $\mathbf{g}_{ij} \leftarrow \text{Linear}(\text{ReLU}(\mathbf{b}_{ij}))$ ▷ $\mathbf{g}_{ij} \in \mathbb{R}^c$
  **return** $\{\mathbf{g}_{ij}\}$
**end function**

**Algorithm S3** Pair denoiser

**Input:** Noisy pair representation $\mathbf{z}$, pair conditioning $\mathbf{g}$

**Output:** Denoised pair representation $\mathbf{z}$

**function** pair_denoiser($\{\mathbf{z}_{ij}\}, \{\mathbf{g}_{ij}\}, N_{\text{block}} = 4, c = 128, c_{\text{mul}} = 128, c_{\text{att}} = 32, N_{\text{head}} = 4$)
  **for** $n \leftarrow 1$ to $N_{\text{block}}$ **do**
    $\{\mathbf{z}_{ij}\} \leftarrow \{\mathbf{z}_{ij}\}$ + TriangleMultiplicationOutgoing($\{\mathbf{z}_{ij}\}$)
    $\{\mathbf{z}_{ij}\} \leftarrow \{\mathbf{z}_{ij}\}$ + TriangleMultiplicationIncoming($\{\mathbf{z}_{ij}\}$)
    $\{\mathbf{z}_{ij}\} \leftarrow \{\mathbf{z}_{ij}\}$ + Conditioning(TriangleAttentionStartingNode, $\{\mathbf{z}_{ij}\}, \{\mathbf{g}_{ij}\}$) ▷ Algorithm S4
    $\{\mathbf{z}_{ij}\} \leftarrow \{\mathbf{z}_{ij}\}$ + Conditioning(TriangleAttentionEndingNode, $\{\mathbf{z}_{ij}\}, \{\mathbf{g}_{ij}\}$)
    $\{\mathbf{z}_{ij}\} \leftarrow \{\mathbf{z}_{ij}\}$ + Conditioning(PairTransition, $\{\mathbf{z}_{ij}\}, \{\mathbf{g}_{ij}\}$)
  **end for**
  **return** $\{\mathbf{z}_{ij}\}$
**end function**

**Algorithm S4** Conditioning

**Input:** Function to be conditioned $\mathcal{F}$, pair representation $\mathbf{z}$, pair conditioning $\mathbf{g}$

**Output:** Pair representation update $\mathbf{u}$

**function** Conditioning($\mathcal{F}, \{\mathbf{z}_{ij}\}, \{\mathbf{g}_{ij}\}$)
  $\mathbf{b}_{ij} \leftarrow \text{AdaLN}(\mathbf{z}_{ij}, \mathbf{g}_{ij})$ ▷ AF3 Algorithm 26 [10]
  $\{\mathbf{b}_{ij}\} \leftarrow \mathcal{F}(\{\mathbf{b}_{ij}\})$
  $\mathbf{u}_{ij} \leftarrow \mathbf{b}_{ij} \odot \text{sigmoid}(\text{Linear}(\mathbf{g}_{ij}), \text{biasinit} = -2.0)$
  **return** $\{\mathbf{u}_{ij}\}$
**end function**

**Algorithm S5** AF2Dock training step

**Input:** Training sample containing bound all-atom complex structure $\mathbf{x}_1$, receptor and ligand all-atom monomer structures $\{\mathbf{x}_{rec,0}, \mathbf{x}_{lig,0}\}$, and their sequences $\{a_{rec}, a_{lig}\}$
  Sample $t \sim \mathcal{U}(0, 1)$
  Split $\{\mathbf{x}_{rec,1}, \mathbf{x}_{lig,1}\} \leftarrow \text{Split}(\mathbf{x}_1)$
  Align $\mathbf{x}_{rec,0} \leftarrow \text{RMSDAlign}(\mathbf{x}_{rec,0}, \mathbf{x}_{rec,1})$
  Align $\mathbf{x}_{lig,0} \leftarrow \text{RMSDAlign}(\mathbf{x}_{lig,0}, \mathbf{x}_{lig,1})$
  Sample $\mathbf{x}_t \leftarrow$ SamplePoseAtT($\mathbf{x}_{rec,0}, \mathbf{x}_{lig,0}, \mathbf{x}_{rec,1}, \mathbf{x}_{lig,1}, t$) ▷ Algorithm S6
  Compute $\{\mathcal{E}_{rec}, \mathcal{E}_{lig}\} \leftarrow \text{ESM}(\{a_{rec}, a_{lig}\})$
  Predict $\hat{\mathbf{x}}_1 \leftarrow \text{AF2Dock}(\mathbf{x}_t, t, \{a_{rec}, a_{lig}\}, \{\mathcal{E}_{rec}, \mathcal{E}_{lig}\})$
  Optimize $\mathcal{L} = \text{AlphaFoldLoss}(\hat{\mathbf{x}}_1, \mathbf{x}_1)$

**Algorithm S6** Sample pose at $t$

**function** SamplePoseAtT($\mathbf{x}_{rec,0}, \mathbf{x}_{lig,0}, \mathbf{x}_{rec,1}, \mathbf{x}_{lig,1}, t, \sigma_{tr} = 30$)
  Sample $\epsilon_{tr} \sim \mathcal{N}(0, \sigma_{tr} I_3)$
  Sample $\epsilon_{rot} \sim \mathcal{U}(SO(3))$
  Interpolate $\mathbf{x}_{rec,t} \leftarrow (1 - t) \cdot \mathbf{x}_{rec,0} + t \cdot \mathbf{x}_{rec,1}$
  Interpolate $\mathbf{x}_{lig,t} \leftarrow (1 - t) \cdot \mathbf{x}_{lig,0} + t \cdot \mathbf{x}_{lig,1}$
  Perturb $\mathbf{x}_{lig,t} \leftarrow \text{Apply}(\mathbf{x}_{lig,t}, (1 - t) \cdot \epsilon_{tr}, (1 - t) \cdot \epsilon_{rot})$
  Combine $\mathbf{x}_t \leftarrow \text{Combine}(\{\mathbf{x}_{rec,t}, \mathbf{x}_{lig,t}\})$
  **return** $\mathbf{x}_t$
**end function**

**Algorithm S7** AF2Dock inference

**Input:** Receptor and ligand all-atom structures $\{\mathbf{x}_{rec,0}, \mathbf{x}_{lig,0}\}$ and their sequences $\{a_{rec}, a_{lig}\}$
**Output:** Predicted all-atom complex structure $\mathbf{x}_1$

Center $\mathbf{x}_{rec,0} \leftarrow \mathbf{x}_{rec,0} - \text{CenterOfMass}(\mathbf{x}_{rec,0})$
Center $\mathbf{x}_{lig,0} \leftarrow \mathbf{x}_{lig,0} - \text{CenterOfMass}(\mathbf{x}_{lig,0})$
Sample $\mathbf{x}_0 \leftarrow \text{SamplePoseAtT}(\mathbf{x}_{rec,0}, \mathbf{x}_{lig,0}, \emptyset, \emptyset, t = 0)$ ▷ Algorithm S6
Compute $\{\mathcal{E}_{rec}, \mathcal{E}_{lig}\} \leftarrow \text{ESM}(\{a_{rec}, a_{lig}\})$
**for** $n \leftarrow 0$ to $N - 1$ **do**
  Let $t \leftarrow n/N$ and $s \leftarrow t + 1/N$
  Predict $\hat{\mathbf{x}}_1 \leftarrow \text{AF2Dock}(\mathbf{x}_t, t, \{a_{rec}, a_{lig}\}, \{\mathcal{E}_{rec}, \mathcal{E}_{lig}\})$
  Interpolate $\mathbf{x}_s \leftarrow \text{InterpolatePoseAtT}(\mathbf{x}_t, \hat{\mathbf{x}}_1, t, s)$ ▷ Algorithm S8
**end for**

**Algorithm S8** Interpolate pose at $t$

**function** InterpolatePoseAtT$(\mathbf{x}_t, \hat{\mathbf{x}}_1, t, s)$
  Align $\hat{\mathbf{x}}_1 \leftarrow \text{RMSDAlignOnReceptor}(\hat{\mathbf{x}}_1, \mathbf{x}_t)$
  Split $\{\mathbf{x}_{rec,t}, \mathbf{x}_{lig,t}\} \leftarrow \text{Split}(\mathbf{x}_t)$
  Split $\{\hat{\mathbf{x}}_{rec,1}, \hat{\mathbf{x}}_{lig,1}\} \leftarrow \text{Split}(\hat{\mathbf{x}}_1)$
  Align $\hat{\mathbf{x}}_{lig,1}, \epsilon_{tr,t}, \epsilon_{rot,t} \leftarrow \text{RMSDAlignReturnTransform}(\hat{\mathbf{x}}_{lig,1}, \mathbf{x}_{lig,t})$
  Interpolate $\mathbf{x}_{rec,s} \leftarrow \frac{(s-t)}{(1-t)} \cdot \hat{\mathbf{x}}_{rec,1} + \frac{(1-s)}{(1-t)} \cdot \mathbf{x}_{rec,t}$
  Interpolate $\mathbf{x}_{lig,s} \leftarrow \frac{(s-t)}{(1-t)} \cdot \hat{\mathbf{x}}_{lig,1} + \frac{(1-s)}{(1-t)} \cdot \mathbf{x}_{lig,t}$
  Perturb $\mathbf{x}_{lig,s} \leftarrow \text{Apply}(\mathbf{x}_{lig,s}, -\frac{(s-t)}{(1-t)} \cdot \epsilon_{tr,t}, -\frac{(s-t)}{(1-t)} \cdot \epsilon_{rot,t})$
  Combine $\mathbf{x}_s \leftarrow \text{Combine}(\{\mathbf{x}_{rec,s}, \mathbf{x}_{lig,s}\})$
  **return** $\mathbf{x}_s$
**end function**

### References


[1] D. Kovtun, M. Akdel, A. Goncearenco, G. Zhou, G. Holt, D. Baugher, D. Lin, Y. Adeshina, T. Castiglione, X. Wang, C. Marquet, M. McPartlon, T. Geffner, E. Rossi, G. Corso, H. Stärk, Z. Carpenter, E. Kucukbenli, M. Bronstein, and L. Naef, bioRxiv (2024).

[2] G. Ahdritz, N. Bouatta, C. Floristean, S. Kadyan, Q. Xia, W. Gerecke, T. J. O'Donnell, D. Berenberg, I. Fisk, N. Zanichelli, B. Zhang, A. Nowaczynski, B. Wang, M. M. Stepniewska-Dziubinska, S. Zhang, A. Ojewole, M. E. Guney, S. Biderman, A. M. Watkins, S. Ra, P. R. Lorenzo, L. Nivon, B. Weitzner, Y.-E. A. Ban, S. Chen, M. Zhang, C. Li, S. L. Song, Y. He, P. K. Sorger, E. Mostaque, Z. Zhang, R. Bonneau, and M. AlQuraishi, Nat. Methods **21**, 1514 (2024).

[3] R. Evans, M. O'Neill, A. Pritzel, N. Antropova, A. Senior, T. Green, A. Žídek, R. Bates, S. Blackwell, J. Yim, O. Ronneberger, S. Bodenstein, M. Zielinski, A. Bridgland, A. Potapenko, A. Cowie, K. Tunyasuvunakool, R. Jain, E. Clancy, P. Kohli, J. Jumper, and D. Hassabis, bioRxiv, 2021.10.04.463034 (2021).

[4] J. Jumper, R. Evans, A. Pritzel, T. Green, M. Figurnov, O. Ronneberger, K. Tunyasuvunakool, R. Bates, A. Žídek, A. Potapenko, A. Bridgland, C. Meyer, S. A. A. Kohl, A. J. Ballard, A. Cowie, B. Romera-Paredes, S. Nikolov, R. Jain, J. Adler, T. Back, S. Petersen, D. Reiman, E. Clancy, M. Zielinski, M. Steinegger, M. Pacholska, T. Berghammer, S. Bodenstein, D. Silver, O. Vinyals, A. W. Senior, K. Kavukcuoglu, P. Kohli, and D. Hassabis, Nature **596**, 583 (2021).

[5] M. Mirdita, K. Schütze, Y. Moriwaki, L. Heo, S. Ovchinnikov, and M. Steinegger, Nat. Methods **19**, 679 (2022).

[6] R. F. Alford, A. Leaver-Fay, J. R. Jeliazkov, M. J. O'Meara, F. P. DiMaio, H. Park, M. V. Shapovalov, P. D. Renfrew, V. K. Mulligan, K. Kappel, J. W. Labonte, M. S. Pacella, R. Bonneau, P. Bradley, R. L.

Dunbrack, Jr, R. Das, D. Baker, B. Kuhlman, T. Kortemme, and J. J. Gray, J. Chem. Theory Comput. **13**, 3031 (2017).

[7] V. Stebliankin, A. Shirali, P. Baral, J. Shi, P. Chapagain, K. Mathee, and G. Narasimhan, Nat. Mach. Intell. **5**, 1042 (2023).

[8] J. P. Roney and S. Ovchinnikov, Phys. Rev. Lett. **129**, 238101 (2022).

[9] C. Mirabello, B. Wallner, B. Nystedt, S. Azinas, and M. Carroni, Nat. Commun. **15**, 8724 (2024).

[10] J. Abramson, J. Adler, J. Dunger, R. Evans, T. Green, A. Pritzel, O. Ronneberger, L. Willmore, A. J. Ballard, J. Bambrick, S. W. Bodenstein, D. A. Evans, C.-C. Hung, M. O'Neill, D. Reiman, K. Tunyasuvunakool, Z. Wu, A. Žemgulytė, E. Arvaniti, C. Beattie, O. Bertolli, A. Bridgland, A. Cherepanov, M. Congreve, A. I. Cowen-Rivers, A. Cowie, M. Figurnov, F. B. Fuchs, H. Gladman, R. Jain, Y. A. Khan, C. M. R. Low, K. Perlin, A. Potapenko, P. Savy, S. Singh, A. Stecula, A. Thillaisundaram, C. Tong, S. Yakneen, E. D. Zhong, M. Zielinski, A. Žídek, V. Bapst, P. Kohli, M. Jaderberg, D. Hassabis, and J. M. Jumper, Nature **630**, 493 (2024).
